# Bicoid-nucleosome competition sets a concentration threshold for transcription constrained by genome replication

**DOI:** 10.1101/2024.12.10.627802

**Authors:** Eleanor A. Degen, Corinne Croslyn, Niall M. Mangan, Shelby A. Blythe

## Abstract

Transcription factors (TFs) regulate gene expression despite constraints from chromatin structure and the cell cycle. Here we examine the concentration-dependent regulation of *hunchback* by the Bicoid morphogen through a combination of quantitative imaging, mathematical modeling and epigenomics in *Drosophila* embryos. By live imaging of MS2 reporters, we find that, following mitosis, the timing of transcriptional activation driven by the *hunchback* P2 (*hb* P2) enhancer directly reflects Bicoid concentration. We build a stochastic model that can explain *in vivo* onset time distributions by accounting for both the competition between Bicoid and nucleosomes at *hb* P2 and a negative influence of DNA replication on transcriptional elongation. Experimental modulation of nucleosome stability alters onset time distributions and the posterior boundary of *hunchback* expression. We conclude that TF-nucleosome competition is the molecular mechanism whereby the Bicoid morphogen gradient specifies the posterior boundary of *hunchback* expression.

## INTRODUCTION

Transcription factors (TFs) bind to sites within cis-regulatory DNA sequences (enhancers) to modulate the expression of associated genes (Levine & Davidson, 2005). How TFs regulate target gene transcription depends on enhancer binding site composition, the contemporaneous TF concentration, as well as competition between DNA binding factors (Zeitlinger, 2020). Theoretical models have highlighted the importance of TF concentration and chromatin context in determining TF binding (Marzen et al., 2013; Mirny, 2010; Narula & Igoshin, 2010). The activity of the TF Bicoid (Bcd) in the *Drosophila* embryo presents an ideal system for studying concentration-sensitive TF-chromatin interactions *in vivo*. Long appreciated as a canonical morphogen, Bcd is expressed in an anterior-posterior (AP) concentration gradient that positions the expression of zygotic segmentation genes early in development (H. Chen et al., 2012; Driever & Nüsslein-Volhard, 1988b, 1988a; Struhl et al., 1989; Wieschaus, 2016; Wolpert, 1994). Target gene expression domains span different ranges of Bcd levels, suggesting that the enhancers mediating these domains differ in their affinity for Bcd (Driever et al., 1989). However, Bcd binding site strength and enhancer affinity fail to predict the Bcd concentration thresholds associated with target gene expression boundaries (Hannon et al., 2017; Ochoa-Espinosa et al., 2005). Prior work has established that the concentration-sensitivity of chromatin accessibility at a locus can explain the degree that Bcd binding to that locus depends on Bcd concentration (Hannon et al., 2017). Yet, we do not fully understand how Bcd-nucleosome competition for occupancy at a locus contributes to precise target gene transcriptional outputs across the gradient.

Maternally-supplied morphogens like Bcd initiate pattern formation despite constraints on transcription imposed by intense cell cycle activity (Briscoe & Small, 2015; Rogers & Schier, 2011). In the syncytial *Drosophila* embryo, 13 synchronous nuclear divisions occur over the first two hours of development. During each mitosis, nuclei export TFs and RNA polymerase II, limiting transcriptional activity. In each of the following interphases, nuclei must rapidly complete DNA replication, reorganize chromatin, and attempt to activate transcription (Blythe & Wieschaus, 2015b, 2016). While intense cellular proliferation characterizes early development, relatively few studies have considered the impact of the cell cycle on transcription prior to large-scale zygotic genome activation. Recent work in early *Drosophila* embryos has shown a coupling between replication and transcription such that inhibition of DNA replication reduces the activity of a Bcd-driven transcriptional reporter (Cho et al., 2022). In addition to TF-chromatin interactions, the influence of the cell cycle on the precise gene expression outputs of the Bcd morphogen gradient has not been fully addressed.

Computational models have built up our understanding of how Bcd regulates patterns of gene expression *in vivo* (Desponds et al., 2016; Eck et al., 2020; Estrada et al., 2016; Fernandes et al., 2022; Lammers et al., 2020; Lopes et al., 2005; Lucas et al., 2018; Segal et al., 2008; Tran et al., 2018). Many studies have focused on Bcd’s activity at *hunchback* P2 (*hb* P2), an enhancer that contributes to the anterior domain of zygotic *hunchback* (*hb*) expression (Struhl et al., 1989). Analyses of static expression measurements have suggested that Bcd binds cooperatively to determine the position and steepness of the anterior *hb* domain’s posterior boundary (Gregor et al., 2007; Park et al., 2019). Bcd’s cooperative binding may require energy-expenditure or a higher order mechanism than pairwise protein-protein interactions (Estrada et al., 2016; Lucas et al., 2018; Park et al., 2019; Tran et al., 2018). Additionally, the TF Zelda and *hb* itself can influence *hb* P2 reporter gene expression (Eck et al., 2020; Fernandes et al., 2022; Park et al., 2019; Simpson-Brose et al., 1994). Models describing transcriptional dynamics measured with the MS2-MCP RNA labeling system have helped detangle the potential regulatory roles of Bcd and additional TFs at *hb* P2 (Eck et al., 2020; Fernandes et al., 2022; Garcia et al., 2013; Lucas et al., 2013, 2018). A recent model suggests that energy-consuming remodeling of chromatin may serve as a mode of Bcd-dependent transcriptional regulation at *hb* P2 (Eck et al., 2020). Modeling *hb* P2-mediated transcriptional dynamics therefore provides a fruitful path toward understanding how *in vivo* context dictates TF concentration-sensitivity.

Here, we define molecular mechanisms by which chromatin and the cell cycle contribute to Bcd concentration-sensitive transcriptional dynamics. We investigate how the initiation of transcription of a highly Bcd-dependent *hbP2-MS2* reporter during nuclear cycle 13 relies on Bcd concentration. By building a stochastic model of transcriptional activation, we propose how Bcd-nucleosome competition and DNA replication determine transcription’s Bcd concentration-sensitivity. We validate the model by testing its predictions through live-imaging and genome-wide sequencing experiments and propose common regulatory mechanisms for Bcd target genes.

## RESULTS

### The activity of the *hunchback* P2 enhancer as a case-study for transcription factor-nucleosome competition

To understand how TF-chromatin interactions influence transcription, we consider the regulation of the zygotic patterning gene *hb*. Early in development, a proximal promoter and pair of enhancers (P2 and shadow) regulate an anterior domain of *hb* expression (Driever & Nüsslein-Volhard, 1989; Perry et al., 2011; Struhl et al., 1989). Later, the distal promoter and the stripe enhancer take over the regulation of *hb*, mediating anterior and posterior stripes of expression (Margolis et al., 1995). As measured by ChIP-seq, Bcd binds to the P2 and shadow enhancers, and to a lesser extent the stripe enhancer (Fig 1A). Genome-wide measurements of Bcd binding and chromatin accessibility suggest that Bcd competes with nucleosomes to bind the *hb* P2 enhancer (Hannon et al., 2017). ChIP-seq measurements in embryos expressing Bcd at low, medium, or high uniform (non-graded) concentrations along the embryo show that Bcd binds to *hb* P2 less at the lower concentration than it does at the higher concentrations (Hannon et al., 2017) (Fig 1A). Chromatin accessibility as measured by ATAC-seq at *hb* P2 also depends on Bcd concentration, suggesting Bcd may reorganize nucleosomes at *hb* P2 (Fig 1B). Using NucleoATAC to predict dyad centers from ATAC-seq fragment size distributions (Schep et al., 2015), we find that two nucleosome dyads are predicted to overlap the *hb* P2 enhancer in embryos mutant for *bcd* (Fig 1B, *bcd^E1^* arrowheads) (Hannon et al., 2017). The *bcd^E1^*mutation represents a complete loss of function and the ATAC measurements therefore reflect the chromatin state observed in the absence of the TF. Increasing the concentration of Bcd gradually reduces the likelihood of dyad prediction suggesting that a high Bcd concentration reduces nucleosome occupancy at *hb* P2 (Fig 1B). The likelihood that a nucleosome occupies a site can also be predicted from the underlying DNA sequence (Segal et al., 2008). A sequence analysis using the Widom-Segal statistical positioning model predicts occupancy of two nucleosomes within *hb* P2 at similar positions to those measured in *bcd^E1^* mutants (Xi et al., 2010) (Fig 1C). These predicted nucleosomes obscure all available Bcd binding sites. A sequence analysis based on the Bcd Position Weight Matrix (PWM) supported by ChIP-Nexus Bcd footprinting (Brennan et al., 2023) reveals nine Bcd binding sites in *hb* P2 overlapping with the observed distal and proximal nucleosomes (Fig 1C & Table 1). Our *in vivo* and *in silico* observations both support a model where Bcd must outcompete nucleosomes in order to access its sites in *hb* P2, where the outcome of Bcd-nucleosome competition depends on Bcd concentration. Therefore, Bcd’s activity at *hb* P2 provides an ideal system for addressing the fundamental problem of how TF-chromatin interactions regulate transcriptional output.

**Figure 1:**
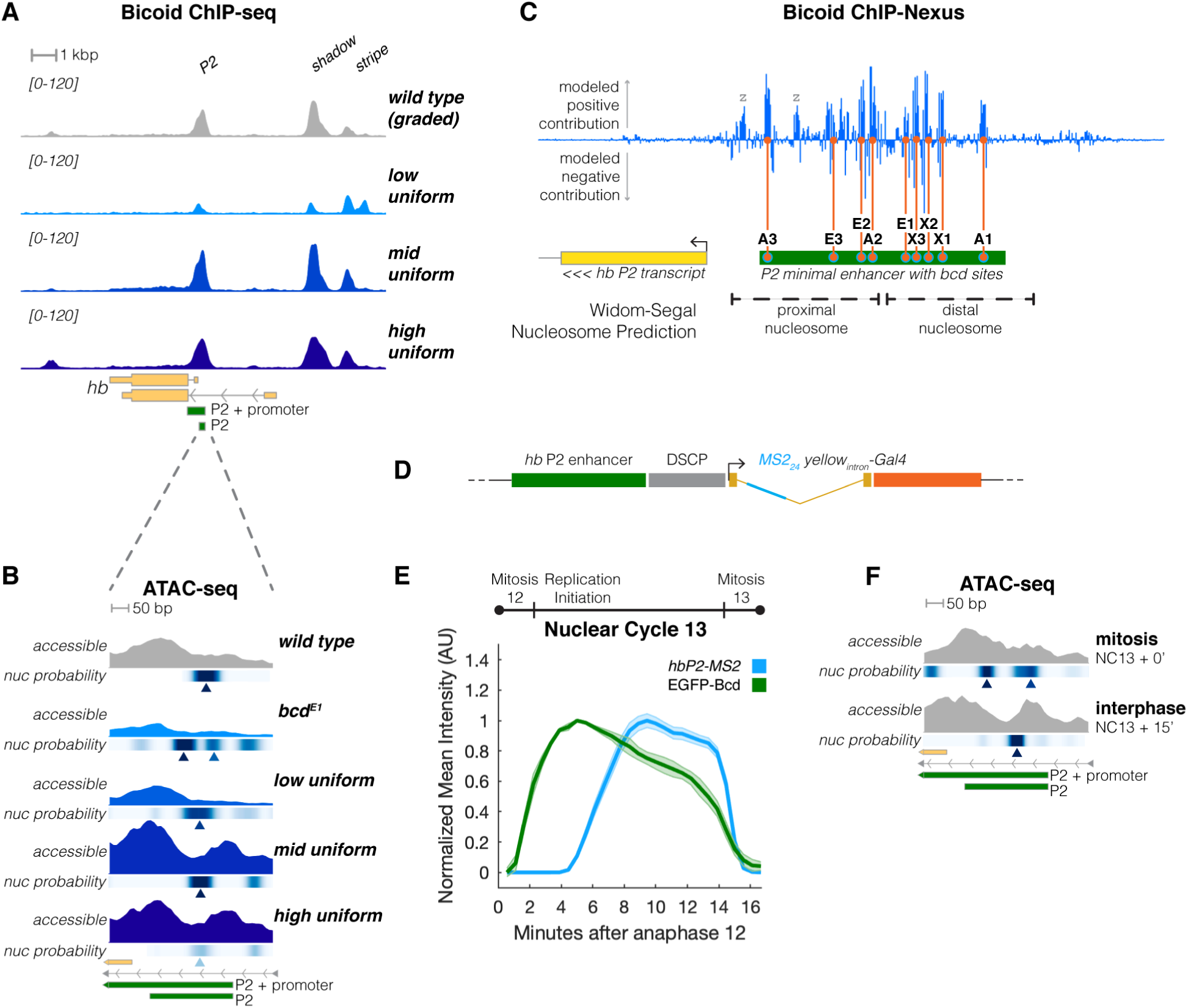
Nucleosome positioning at the *hb* P2 locus is sensitive to Bicoid concentration and cell cycle timing. **A)** Bcd binds in a concentration-dependent manner to *hb* regulatory elements. Shown is a 15 kb region flanking the *hb* locus including the P2, shadow, and stripe regulatory elements (indicated at top), and counts-per-million (CPM) normalized ChIP-seq for Bcd protein. Mean binding measurements for wild type embryos (top row) is shown in comparison to three mutant lines that express Bcd uniformly across the AP axis at low, medium, or high concentrations. Data from three independent biological replicates were averaged for each genotype. Plotted ChIP-seq coverage ranges from 0-120 CPM for all conditions shown. The genomic locations of the minimal P2 enhancer and extended P2+promoter regions are shown in green (bottom). Scale bar = 1 kb. **B)** Bcd is required to establish accessible chromatin at the P2 enhancer element. Shown is a 0.6 kb detail of the region upstream of the transcription start site of the *hb* P2 isoform including the entire minimal P2 enhancer element (green, bottom). ATAC-seq data corresponding to accessible regions and nucleosome dyad positions predicted by NucleoATAC (nuc probability) are shown for five genotypes: wild type, *bcd^E1^*, and low, medium, and high uniform Bcd. Predicted dyad positions referred to in the text are highlighted with arrowheads. Data from three independent biological replicates were averaged for each genotype. Plotted open ATAC coverage ranges from 0-20 CPM for all conditions shown. Scale bar = 50 bp. **C)** ChIP-Nexus data supports the presence of at least nine Bcd binding sites in the minimal P2 enhancer. Shown is the same genomic region as in (B). Positive and negative contributions to Bcd binding are plotted (blue, Brennan, et al. 2023) and the positions are shown of six binding sites identified through DNase footprinting (A1-3, X1-3), in addition to three <u>e</u>xtra motifs with PWM scores greater than 80% (E1-3). Two footprints are observed overlapping motifs for Zld (“z”). It is unclear if this represents Bcd binding at these sites, or footprinting of co-immunoprecipitated Zld proteins. The lower portion of the panel shows the positions of two nucleosomes predicted by NuPoP, and a schematic of the P2 enhancer with nine Bcd binding sites. **D)** Schematic of a minimal *hbP2-MS2* reporter: the minimal *hb* P2 enhancer (green) drives the expression of a chimeric *MS2(24)-yellow intron -Gal4* reporter gene through the heterologous Drosophila Synthetic Core Promoter (DSCP, grey). Sequences encoding 24 MS2(v5) hairpins (blue) were introduced to the 5’ end of the *yellow* intron (yellow) upstream of the *gal4* coding sequence (orange). **E)** Expression of *hbP2-MS2* initiates after nuclear import of Bcd and expected replication initiation. Shown is the mean reporter activity of *hbP2-MS2* (blue) and mean nuclear intensity of EGFP-Bcd (green) measured over Nuclear Cycle 13 (NC13). All three measurements were performed in separate embryos. The timeline (top) highlights the cell cycle landmarks relative to the period of NC13, timed from the beginning of anaphase 12. **F)** Nucleosome occupancy over the P2 element fluctuates over the cell cycle. ATAC data over the same region as in (B) is plotted, but for wild type embryos collected at mitotic metaphase (top, NC13+0’) and late interphase (bottom, NC13 + 15’, Blythe and Wieschaus, 2016). Plotted ATAC coverage ranges from 0-20 CPM for all conditions shown. Scale bar = 50 bp.

**Table 1:** Bcd binding sites within *hb* P2. • **Motif Name:** The name of the motif as described in Figure 1 and accompanying text. • **Strand (ref):** The strand (+ or -) of motif occurrence relative to the reference genome. • **Sequence:** The sequence of the motif, delimited by bases predicted by ChIP-Nexus to contribute positively to Bcd binding. The sequences are aligned by the frequent common core ATCC bases indicated in bold. • **Coordinates (dm6):** The genomic coordinates in the dm6 reference genome assembly for the listed sequence.

| Motif Name | Strand (ref) | Sequence | Coordinates (dm6) |
| --- | --- | --- | --- |
| A1 | - | CGTA**ATCC** | chr3R:8694875-8694882 |
| X1 | - | TA**AGCTG** | chr3R:8694832-8694838 |
| X2 | - | TA**AGCTC** | chr3R:8694816-8694822 |
| X3 | + | TG**ATCCG** | chr3R:8694808-8694814 |
| E1 | - | AA**ATCC** | chr3R:8694799-8694804 |
| A2 | - | CTCTA**ATCC** | chr3R:8694765-8694773 |
| E2 | + | TG**ATCCA** | chr3R:8694754-8694760 |
| E3 | - | CCTCA**ATCCG** | chr3R:8694734-8694743 |
| A3 | - | CATCTA**ATCC** | chr3R:8694660-8694669 |

### Live transcriptional reporters allow for the measurement of stages of the transcriptional cycle regulated by upstream transcription factor activity

To read out the effects of Bcd-nucleosome competition on transcription, we created an MS2 reporter for the *hb* P2 enhancer (*hbP2-MS2*). *hb* P2 and a *Drosophila* synthetic core promoter in our reporter drive the expression of a series of 24 MS2 stem loops, facilitating live imaging with the MS2-MCP system (Garcia et al., 2013; Lucas et al., 2013) (Fig. 1D). This reporter does not contain the *hb* P2 promoter nor any additional genomic sequence from the *hb* locus, allowing us to directly evaluate the impact of Bcd’s binding to its sites in *hb* P2 and minimize regulation by additional factors (Fig. 1D). Importantly, NuPoP predicts that a pair of nucleosomes overlap the P2 enhancer within the reporter construct at similar positions to the nucleosomes predicted in the genomic context. While prior studies have analyzed the expression patterns of *hb* P2 MS2 reporters, these reporters reflected the combined activities of *hb* P2 and the P2 promoter (Eck et al., 2020; Fernandes et al., 2022; Garcia et al., 2013; Lucas et al., 2013, 2018). We measured *hbP2-MS2* expression during nuclear cycle 13 (NC13), an on-average 20-minute syncytial cell cycle bounded by near-synchronous nuclear divisions. Beyond minimizing cell cycle variance between nuclei, NC13 precedes large-scale zygotic genome activation and minimizes regulation of *hb* P2 by secondary zygotic factors. Therefore, *hbP2-MS2* dynamics in NC13 can primarily reflect the effects of maternally supplied factors such as Bcd. Quantification of *hbP2-MS2* expression across NC13 reveals average reporter transcription is delayed until around five min into NC13, after which mean reporter activity increases linearly until reaching a maximum at around 10 min into the nuclear cycle (Fig. 1E, blue). Similar patterns of transcription have been produced by *hb* P2+P2 promoter reporters (Garcia et al., 2013; Lucas et al., 2013). We aimed to evaluate how *hbP2-MS2* expression depends on Bcd concentration.

Cumulative transcriptional output of a gene expressed during late embryonic cleavage stages depends on features of transcriptional dynamics, including how soon after mitosis nuclei initiate transcription, the rate of RNA polymerase II (Pol II) loading onto the gene, Pol II bursting dynamics at the promoter, and the number of nuclei that activate transcription. To test the hypothesis that a feature of transcription reflects Bcd’s concentration-sensitive role at *hb* P2, we characterized our *hbP2-MS2* reporter’s activity by live-imaging a series of 19 embryos over NC13. We measured the transcriptional dynamics of 1,904 individual nuclei spanning the anterior 60% of the embryo, with 1,384 of these nuclei expressing the reporter (Fig 2A). We extracted features of the dynamics, including the time at which an MCP-GFP spot appears within a nucleus (the “transcriptional onset time” of that nucleus, (Fig 2B)), as well as the rate of increase in MCP-GFP spot fluorescence over the first minute following the transcriptional onset of a nucleus (the “Pol II loading rate” of that nucleus (Fig 2B)). Additionally, we quantified the fraction of nuclei at Bcd concentrations along the anterior-posterior axis (AP position) that initiate transcription in NC13 (the “fraction of active nuclei”). A heatmap representation of per-nucleus MS2 signal indicates nuclei vary in the time when they first initiate reporter transcription (Fig 2A). We hypothesized that one or more of the features of transcriptional dynamics – transcriptional onset time, Pol II loading rate, and fraction of active nuclei – would correlate with the change in Bcd concentration across the AP axis.

**Figure 2:**
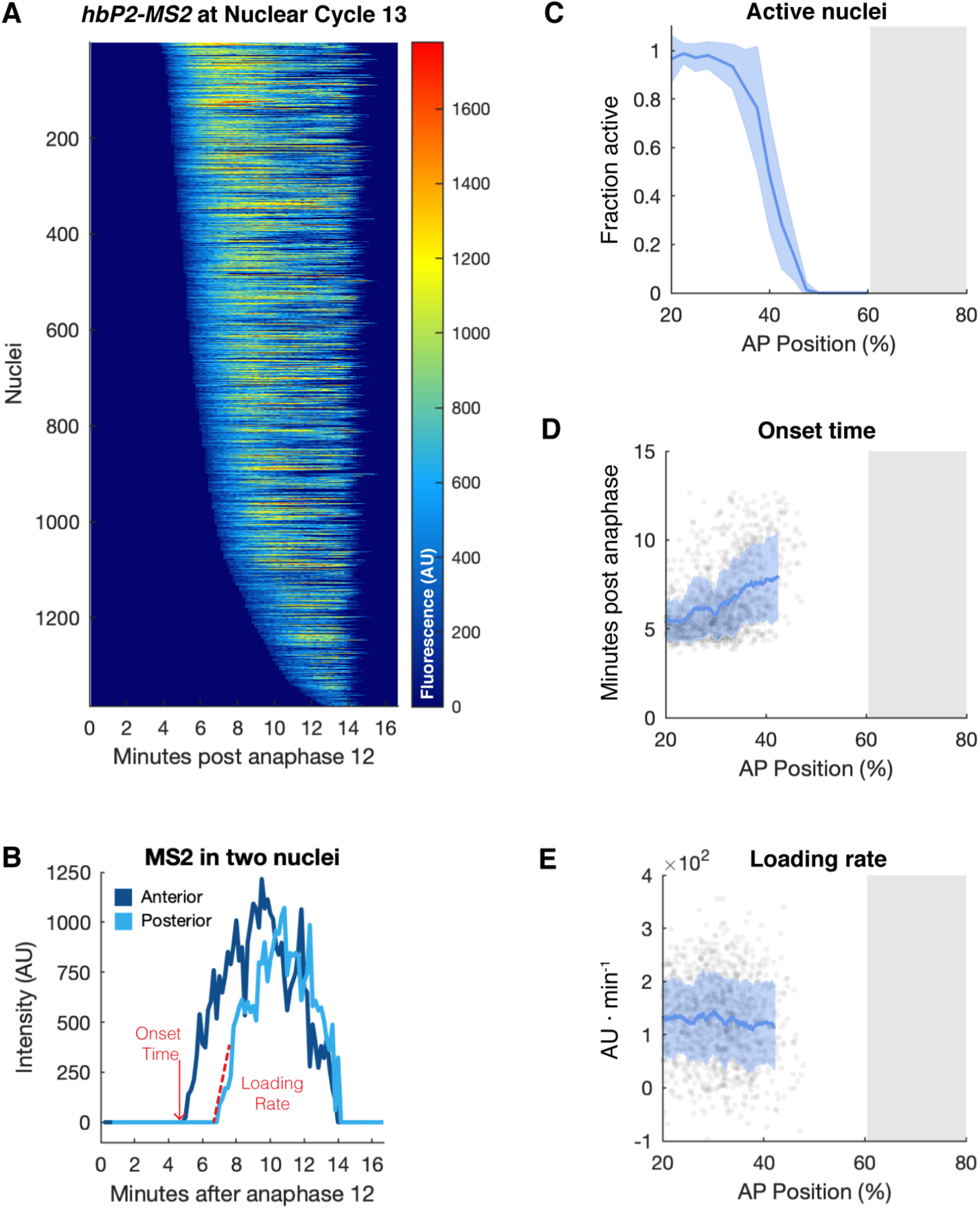
Transcriptional onset time of the *hbP2-MS2* reporter varies as a function of AP position. **A)** A heatmap of *hbP2-MS2*::MCP-GFP fluorescence intensity profiles for ∼1400 individual nuclei (n = 19 independent movies) over NC13 is shown. The x-axis corresponds to minutes following anaphase 12, normalized to 20 minutes. The y-axis represents individual nuclei measured and scored as positive for focal expression of MCP-GFP, and sorted by scored onset time. The fluorescence intensity is depicted according to the colorbar on the right. **B)** Shown are the MS2::MCP-GFP measurements for two representative nuclei, one from the anterior (dark blue) and another from the posterior (light blue) of the *hb* P2 expression domain. Indicated in red are the portions of the data that yield the onset time and loading rate measurements. While onset times show high variance across the *hb* P2 expression domain, loading rates and maximal intensities show less variation. **C)** The fraction of nuclei actively transcribing the reporter does not vary over the region of greatest change in Bcd concentration. The fraction of active nuclei per 2.5% AP bin was calculated and the mean value between movies was plotted (blue ± standard deviation). The gray box in this plot and the following from 60-80% AP position indicates that no movies were taken within this region. **D)** Reporter onset times show an inverse correlation with Bcd concentration. The onset time of *hbP2-MS2* expression was calculated and plotted as a function of AP position (gray points). A rolling average of onset times (n = 100) is shown (blue ± standard deviation) across the 95% C.I. of active nuclei AP positions. **E)** The average Pol II loading rate of active nuclei does not vary with Bcd concentration. The loading rate of *hbP2-MS2* was calculated and plotted as a function of AP position (gray points). A rolling average of loading rates (n = 100) is shown (blue ± standard deviation) across the 95% C.I. of active nuclei AP positions.

### The timing of transcriptional onset following mitosis reads out changes in Bicoid concentration

Out of the three features of transcriptional dynamics, we find that *hbP2-MS2* transcriptional onset time following mitotic exit (“onset time”, hereafter) exhibits the highest sensitivity to Bcd concentration (Fig. 2C-E). In contrast, the fraction of active nuclei is not highly sensitive to Bcd concentration, as it plateaus in the anterior of the embryo; although Bcd concentrations drop by ∼55% between 20 and 35% egg length (EL), almost 100% of nuclei initiate transcription across these AP positions (Fig. 2C). Similarly, the mean Pol II loading rate does not change substantially across the *hbP2-MS2* expression domain and displays a low sensitivity to Bcd concentration (Fig. 2E). This finding fits with prior observations of little correlation between the transcription rates and gene expression boundaries of Bcd targets (Fukaya, 2021). Mean onset time, however, correlates more with AP position, supporting this feature’s higher sensitivity to changes in Bcd concentration (Fig. 2C). If onset times are indeed sensitive to Bcd concentration, then altering the concentration of Bcd should have a corresponding effect on onset time distribution. To test the effects of changing Bcd concentration, we generated mutant embryos that express Bcd uniformly (Hannon et al., 2017). Comparison of the average fluorescence intensity of uniform EGFP-Bcd (tub>uEGFP-Bcd) to wild type (graded) reveals that the uniform line expresses Bcd at a concentration equivalent to the Bcd level at 38% EL in wild type (Fig. 3A). By *in situ* hybridization, *hbP2-MS2* is uniformly expressed in embryos expressing this level of uniform Bcd (Fig 3B). By live imaging, onset times are also uniform and overlap with the wild type distribution at 37% EL, closely matching the 38% EL overlap position of the graded and uniform Bcd concentration profiles (Fig. 3C). As changes in onset times precisely follow changes in Bcd concentration, we conclude onset time likely reflects the direct consequences of Bcd-chromatin interactions on transcription.

**Figure 3:**
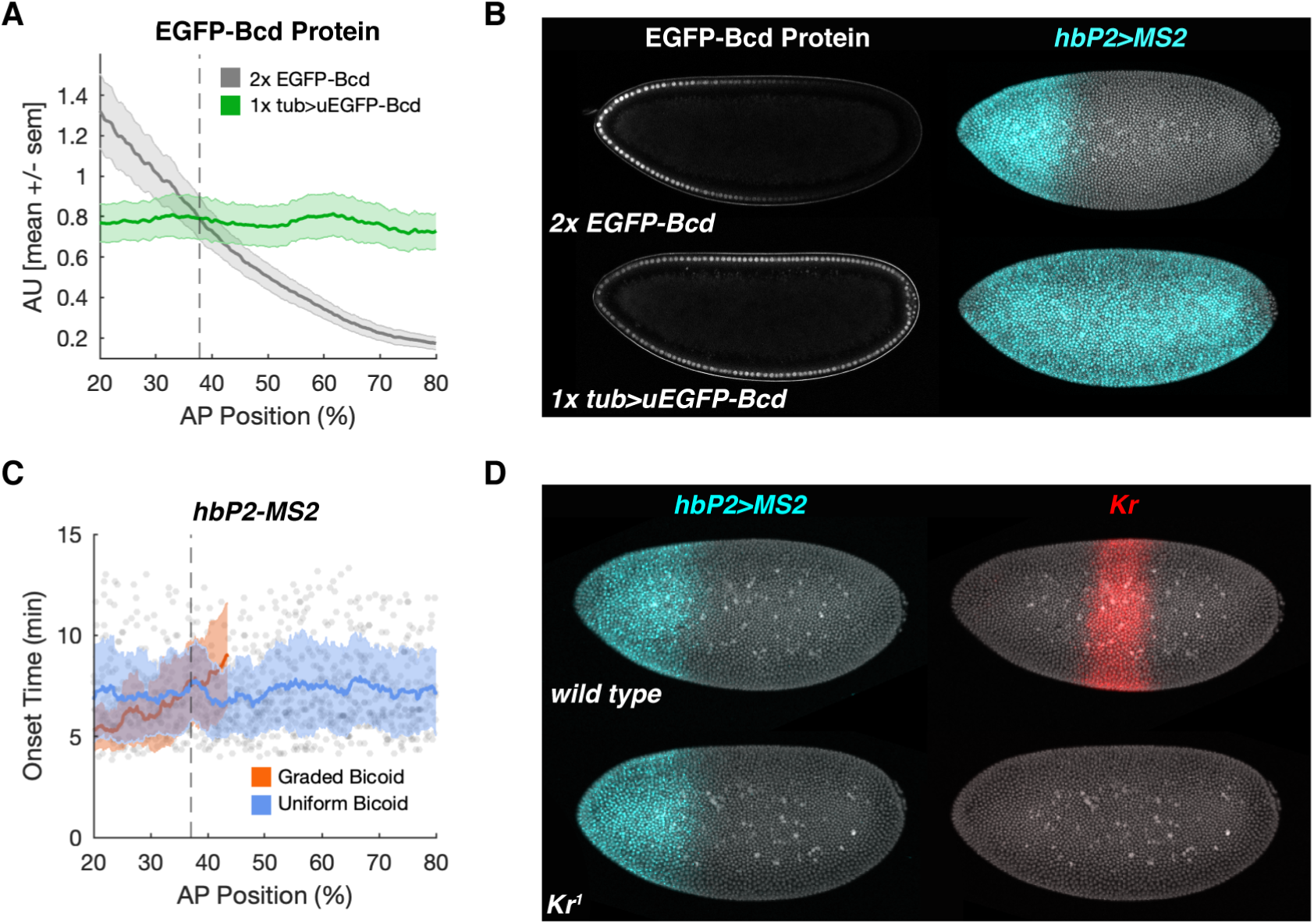
Transcriptional onset time of the *hbP2-MS2* reporter is a direct reflection of Bicoid concentration. **A)** High levels of uniformly expressed Bcd produce uniform Bcd concentrations typically observed at ∼38% egg length. Wild type, graded EGFP-Bcd (2x EGFP-Bcd, n = 6 embryos, gray) or uniform EGFP-Bcd (1x tub>uEGFP-Bcd, n = 8 embryos, green) was live-imaged and nuclear fluorescence intensity during NC13 was quantified and plotted as a function of AP position. A rolling average of fluorescence within a 10% AP window is shown ± standard error of the mean. The dotted line indicates the approximate AP position where uniform Bcd expression coincides with graded Bcd (38%). **B)** Uniform Bicoid drives uniform expression of *hbP2-MS2* across the entire AP axis. At left is shown EGFP-Bcd fluorescence (gray) in representative live embryos expressing either 2x EGFP-Bcd (top) or 1x tub>uEGFP-Bcd (bottom). EGFP emissions from a midsagittal confocal section over the last three minutes of NC13 interphase were summed to produce the images shown. Shown at right are hybridization chain reaction images for *hbP2-MS2-Gal4* (cyan) in either 2xEGFP-Bcd (top) or 1x tub>uEGFP-Bcd (bottom) fixed embryos. Nuclear DNA (DAPI) is shown in gray. The *hbP2-MS2-Gal4* signal extends across the entire AP axis in embryos expressing uniform Bicoid. **C)** Flattening the Bcd gradient flattens the onset time distributions of *hbP2-MS2* to reflect the mean onset time observed at 37% EL. Shown are onset time distributions for *hbP2-MS2* measured in uniform Bcd (blue) and graded Bcd (red) as described in panel A. The vertical dashed line indicates the computed AP axis position where the uniform matches the distribution of graded Bcd *hbP2-MS2* onset times. Plotted gray points indicate individual measurements from the uniform Bcd experiment only, lines represent rolling averages of onset times (n = 50 ± standard deviation) over the 95% C.I. of AP positions with active nuclei. **D)** Kr does not regulate the posterior boundary of *hbP2-MS2*. Gene expression in embryos from a cross between *hbP2-MS2-gal4 Kr^1^/SM6a* individuals was measured by hybridization chain reaction. Left panels show representative *hbP2-MS2* reporter activity (cyan) in a wild type (*Kr*/+) embryo (top) compared with reporter activity in a homozygous *Kr* mutant embryo (bottom), both staged at early NC14. Right panels show *Kr* transcript expression (red) in the same embryos pictured at left. The *Kr*^1^ allele is RNA null. DAPI staining is shown (gray) in both panels to facilitate visualization of the specimens.

### Transcriptional repressors do not contribute to the *hunchback* P2-mediated expression domain

We also ruled out the possibility that repressive TFs influence the delay in transcriptional onset at the boundary of the *hbP2-MS2* domain. While it has been shown that the posterior boundaries of several Bcd targets are shaped by opposing repressor gradients (H. Chen et al., 2012), it is likely that the posterior boundary of *hb* P2 is set directly by a Bcd concentration threshold. To confirm *hbP2-MS2* onset times can read out the direct effects of changing Bcd concentrations at *hb* P2, we sought to rule out possible repressive feedback from maternal and zygotic factors in the patterning gene network. We assert that Capicua (Cic), Runt (Run), Knirps (Kni), and Krüppel (Kr), repressive TFs expressed in domains either overlapping or bordering anterior *hb* expression, do not contribute to *hb* P2 activity. *cic* mutants impact *hb* expression associated with the stripe regulatory element and not the early *hb* expression related to the P2/shadow enhancers (Löhr et al., 2009). Furthermore, Cic does not bind to the *hb* locus (Keenan et al., 2020). In addition, *run* mutants do not affect the *hb* P2 expression domain (H. Chen et al., 2012). *kni* and *Kr* mutants also do not significantly affect early *hb* expression that corresponds to P2 promoter activity (Haroush et al., 2023). While the *hb* P2 enhancer does not contain PWM matches to the Kni motif, we find a potential Kr binding site that overlaps the A3 Bcd motif on the opposite strand (Fig. S1). Kr is a gap gene expressed in a stripe at the posterior boundary of the anterior *hb* domain, suggesting it could refine the posterior boundary of *hb* P2-mediated expression. To confirm that Kr does not regulate *hbP2-MS2* expression, we measured the *hbP2-MS2* reporter in *Kr^1^* mutant embryos. Loss of *Kr* does not change the position of *hbP2-MS2* expression, demonstrating that transcriptional repression by Kr does not limit the posterior boundary of the *hb* P2 domain (Fig. 3D). Therefore, the Bcd concentration gradient likely serves directly as the primary source of positional information for *hb* P2-mediated transcription.

**Supplemental Figure S1, related to Figure 3:**
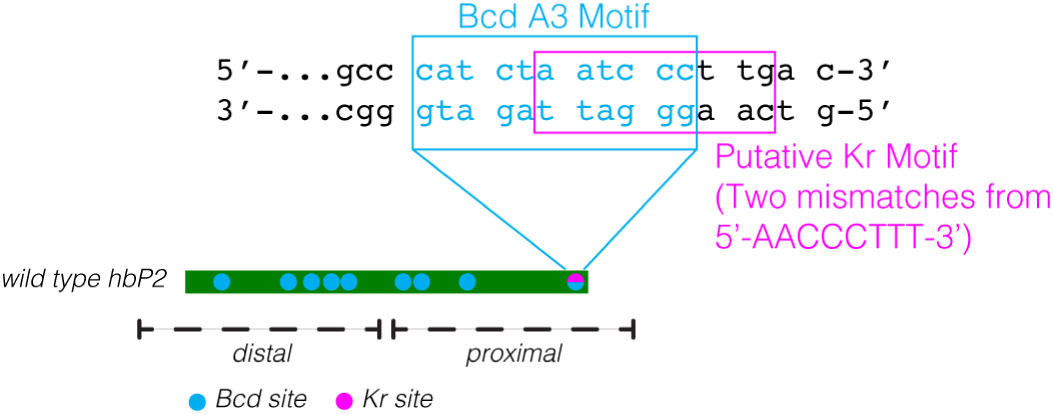
A Kr motif in *hb* P2. Scanning the *hb* P2 sequence with a Kr binding site position weight matrix, one degenerate Kr motif is identified at a match cutoff of ≥85%. This motif overlaps with the Bcd A3 motif at the 3’ end of the *hb* P2 sequence.

### Modeling the determination of transcriptional onset times

Transcription in early development takes place during a time of intense proliferation, where both TF-nucleosome competition and features of the cell cycle can impact gene regulation. Nuclei export TFs and Pol II during mitosis and then reimport the factors in the following interphase, exemplified by the nuclear EGFP-Bcd concentration dynamics during NC13 (Fig. 1E, green). Accessibility at *hb* P2 increases between mitosis 12 and the interphase of NC13, potentially reflecting Bcd’s competition with nucleosomes as nuclear Bcd levels rise (Fig. 1F). Genome replication following mitosis could also influence the transcriptional process, as transcription in syncytial stage *Drosophila* embryos requires the competition of DNA replication (Cho et al., 2022). We developed a mathematical model to investigate how these components of the dynamic genomic environment contribute to the Bcd concentration-sensitive expression of the *hbP2-MS2* reporter.

The sensitivity of *hbP2-MS2* onset time to Bcd concentration implies that Bcd concentration determines the rate at which the reporter’s promoter shifts from a transcriptionally inactive to an active state. We therefore built a two-state model of promoter activity that can simulate transcriptional onset times (Fig. 4A). In this model, following mitosis, the promoter of the reporter starts the nuclear cycle in an “OFF” state where transcription cannot occur. As the nuclear cycle progresses, the promoter can transition to an “ON” state where Pol II transcribes the reporter. To account for the Bcd concentration-sensitivity of the timing of transcriptional onset, we defined the rate at which a nucleus transitions from OFF to ON, *k_on_*, as a function of Bcd concentration (Fig. 4A). The simple two-state, one-transition structure of the model isolates the role of Bcd concentration in transcriptional activation. In the model, Bcd binding at *hb* P2 leads to an increase in *k_on_* through a mechanism described by *k_on_*(*[Bcd]*(*x*,*t*)). By simulating onset times with different forms of *k_on_*(*[Bcd]*(*x*,*t*)), we tested hypotheses about how the *hb* P2 enhancer governs the Bcd-concentration sensitivity of *hbP2-MS2* transcription. In the following, we assess whether different definitions of *k_on_*—including a Michaelis-Menten formulation, Hill function, and model with competitive nucleosome binding—allow the model to reproduce the measured *hbP2-MS2* transcriptional onset time distribution. We first sought to determine whether Bcd-nucleosome competition alone regulates the *hbP2-MS2* expression domain. We employ a version of Gillespie’s stochastic simulation algorithm to simulate the OFF-ON transition in individual nuclei (see Supplemental Information for details) and then compare simulated onset time measurements to the *in vivo* data.

**Figure 4:**
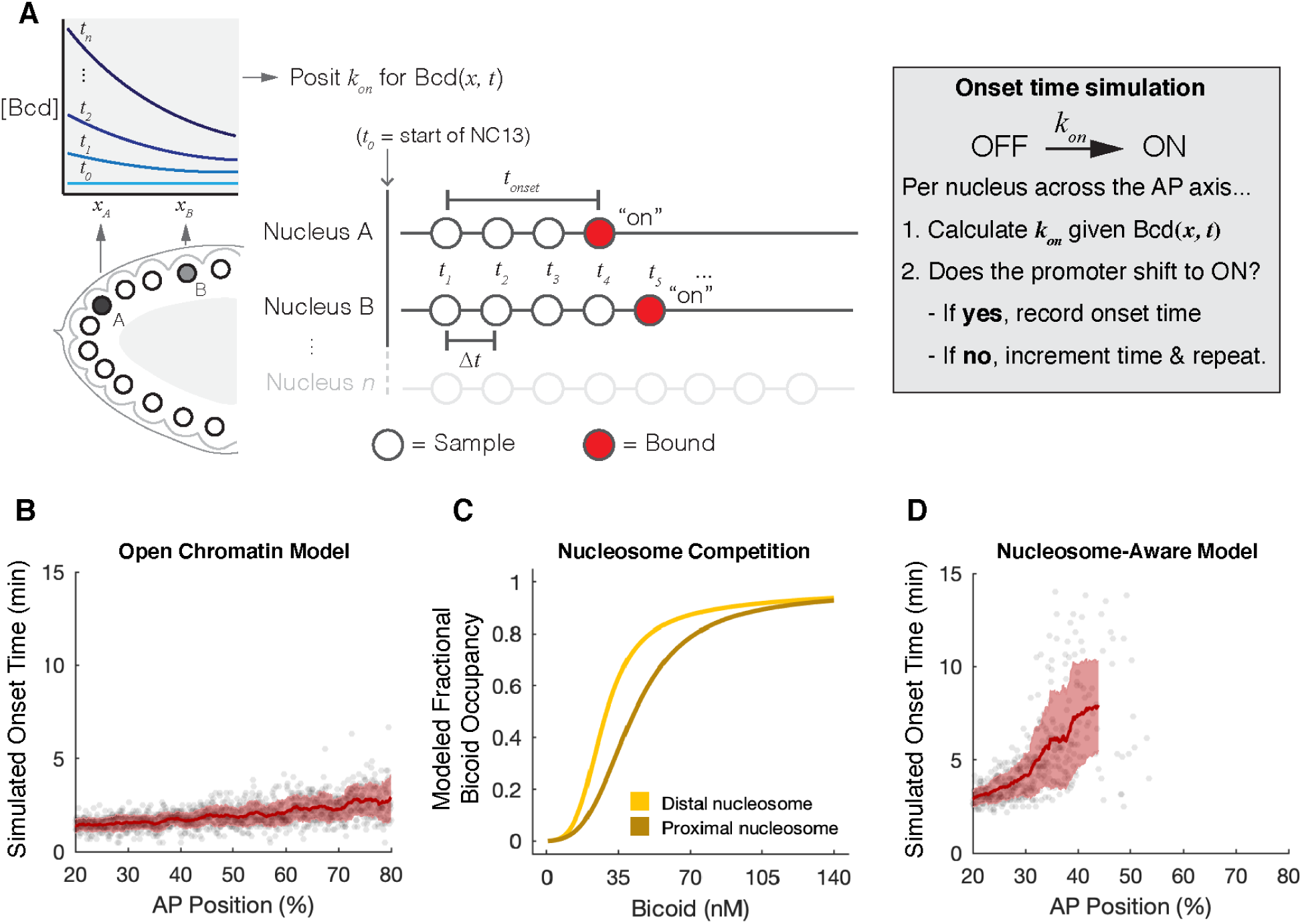
A nucleosome-aware stochastic model for onset time prediction produces a posterior boundary for simulated *hbP2-MS2* activity. **A)** Cartoon schematic of the initial simulation strategy demonstrated for two example nuclei at NC13. Two nuclei (A & B, lower left) in the embryo encounter both spatial and temporal differences in Bcd concentration throughout the cell cycle (top left), which can be expressed as Bcd(*x, t*). The transition to an “on” state is defined by *k_on_* and in this simulation depends solely on whether Bcd binds successfully to the target. Beginning at anaphase of NC12, the value of Bcd(*x, t*) is evaluated and *k_on_* is posited for each nucleus at timepoint *t_i_* until a nucleus is scored as “on”. If, at timepoint *t, k_on_* yields a wait-time for binding less than the time-step (*Δt*), the nucleus is scored as ‘on’ at time *t* (*t_onset_*). If the calculated wait-time is greater than *Δt*, the timepoint is incremented to *t+Δt* and the process repeats. This process yields a *t_onset_* for each nucleus in the field *x*. **B)** Modeling Bcd binding with an open chromatin model reflecting Michaelis-Menten kinetics fails to produce spatiotemporal onset time patterns seen *in vivo*. All plotted data are modeled onset times. Gray points represent individual modeled onset times. The red trace is the rolling mean ± standard deviation of modeled onset times over the 95% C.I. of AP positions with modeled active nuclei. **C)** Over its physiological range of expression, Bcd is predicted to compete effectively with nucleosomes for occupancy of *hb* P2 DNA. Shown is the calculated fractional Bcd occupancy predicted by a TF-nucleosome competition model over the region covered by the proximal (brown) and distal (yellow) nucleosomes. The TF-nucleosome competition model was applied using *K_O_* = 9 nM, *K_N_* = 2610 nM, and *L* = 700 for each nucleosome, and using *n* = 4 and *n* = 5 for the proximal and distal nucleosomes respectively. **D)** Invoking a nucleosome-aware model for Bcd binding to *hb* P2 predicts several features of the *in vivo* onset time distributions. This model captures the distinct posterior boundary to *hb* P2 expression. All plotted data are modeled onset times. Gray points represent individual modeled onset times. The red trace is the rolling mean ± standard deviation of modeled onset times over the 95% C.I. of AP positions with modeled active nuclei. While the mean and variance of simulated onset times trend towards *in vivo* measurements, there are notable deviations. This simulation was performed using *K_O_* = 9 nM, *K_N_* = 2610 nM, *L* = 700, and *V_max_* = 1.

### Transcription factor-nucleosome competition allows the model to predict a Bicoid concentration threshold

We assessed whether chromatin accessibility influences the Bcd concentration-sensitivity of *hbP2-MS2* first by ruling-out that an interaction between Bcd and open chromatin could produce the observed onset time distribution. To simulate Bcd binding non-cooperatively to open chromatin, we defined *k_on_* according to Michaelis-Menten reaction kinetics (Supplemental Information). We find that, with this definition of *k_on_*, the model produces poor onset time predictions. The model notably fails to predict a posterior boundary of expression and predicts transcriptional onset substantially prior to the 4.5-minute onset time minimum observed in the *in vivo* data (Fig. 4B, Supplemental Information). Therefore, beyond binding open chromatin, Bcd likely encounters barriers to transcriptional activation that delay onsets times and prevent *hbP2-MS2* expression at low Bcd levels. As we expect no repressive contribution from other maternal or zygotic factors, we propose the nucleosomes at *hb* P2 provide a barrier to transcriptional activation that delays the onset of transcription. While nucleosomes likely hinder Bcd binding to *hb* P2, Bcd binding to open chromatin through a pairwise cooperative mechanism could theoretically also produce a posterior boundary of target gene expression. We discuss the insufficiency of Hill-like cooperative binding on naked chromatin in the context of our two-state model in the Supplemental Information.

Given the failures of the open chromatin models, we incorporated a previously described model for TF-nucleosome competition into the definition of *k_on_* (Mirny, 2010). Mirny’s TF-nucleosome competition model takes the form of a Monod-Wyman-Changeux model of allostery (Mirny, 2010; Monod et al., 1965). To estimate TF and nucleosome occupancies across different TF concentrations, the TF-nucleosome competition model requires a short list of biophysical parameters: a TF’s affinity for its sites in the absence of a nucleosome (*K_O_*), a TF’s affinity for its sites in the presence of a nucleosome (*K_N_*), a nucleosome stability parameter (*L*), and the number of TF binding sites (*n*). We modeled Bcd’s competition with the proximal and distal nucleosomes using their respective numbers of Bcd binding sites (*n* = 4 and *n* = 5, Fig 1C) and left *K_O_*, *K_N_*, and *L* as free parameters that describe Bcd’s competition with both nucleosomes. To estimate the Bcd occupancy per site at the proximal (*Y_prox_*) and distal (*Y_dist_*) nucleosomes, we applied the TF-nucleosome competition model,

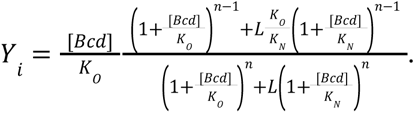

To incorporate the influence of nucleosomes into the two-state model, we set *k_on_* as proportional to the probability Bcd evicts both nucleosomes, *Y_dist_*\* *Y_prox_*:

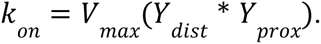

Defining *k_on_* according to Bcd-nucleosome competition allows the two-state model of transcriptional activation to predict a posterior boundary of expression (Fig. 4D). We estimated best-fit values for the parameters *K_O_, K_N_* and *L* by sweeping across ranges of possible values and evaluating simulation outputs (see Supplemental Information for details). We find the best fit parameters fall within the ranges of prior measurements or reasonable estimates, and successfully simulate a posterior expression boundary that aligns with *in vivo* observations (Fig. 4D, Supplemental Information). The combination of *K_O_* = 9 nM, *K_N_* = 2610 nM and *L* = 700 predicts the posterior boundary position of our *hbP2-MS2* measurements very well, deviating by only 0.26% EL (Fig. 4D). *K_O_* = 9 nM falls within the range of published estimates of Bcd’s affinity for its sites *in vitro* (Burz et al., 1998; Burz & Hanes, 2001; Hannon et al., 2017; Ma et al., 1996). *K_N_* = 2610 nM and *L* = 700 are within the ranges approximated in previous work, specifying *K_O_*/*K_N_*= 0.1-0.001 and *L* = 100-1000 (Mirny, 2010). As the nucleosome-dependent model predicts the data’s posterior boundary with reasonable parameter estimates, these results support the hypothesis that nucleosomes play a regulatory role at *hb* P2.

### Nucleosome stability modulates the Bicoid concentration threshold

We tested the conclusion that Bcd-nucleosome competition sets a concentration threshold for transcription by measuring and then modeling the *hbP2-MS2* expression boundary in conditions of altered nucleosome stability. To alter nucleosome stability, we manipulated the activity of the TF Zelda (Zld) at the reporter, as Zld can disrupt nucleosomes and increase chromatin accessibility upon binding to a locus (Liang et al., 2008; Sun et al., 2015). Prior ATAC-seq data shows a slight reduction in chromatin accessibility at the *hb* P2 enhancer in *zld* germline clone mutants (Fig. 5A). Furthermore, the *hb* P2 enhancer contains a single Zld motif occluded by a NuPoP-predicted nucleosome (Fig. 5B). We, therefore, propose that Zld indirectly contributes to *hb* P2-mediated transcription by influencing nucleosome stability and the dynamics of Bcd-nucleosome competition. We first tested the effects of nucleosome stabilization by characterizing *hbP2-MS2* expression in Zld knockdown embryos. We find that, in *zld*-RNAi embryos, the posterior boundary of the fraction of *hbP2-MS2* active nuclei is shifted slightly toward the anterior (Fig. 5C, dark blue). We next tested the effects of nucleosome destabilization by increasing the recruitment of Zld to the *hbP2-MS2* reporter by adding a Zld motif to *hb* P2, thereby creating a [*hbP2* + *1xZld*] *MS2* reporter (Fig. 5B). As indicated by the fraction of active nuclei, we find that [*hbP2* + *1xZld*] *MS2* is expressed farther to the posterior than the wildtype *hbP2-MS2* reporter (Fig. 5C, green). Our experimental results point to a correlation between nucleosome stability and the Bcd concentration threshold required for transcription.

**Figure 5:**
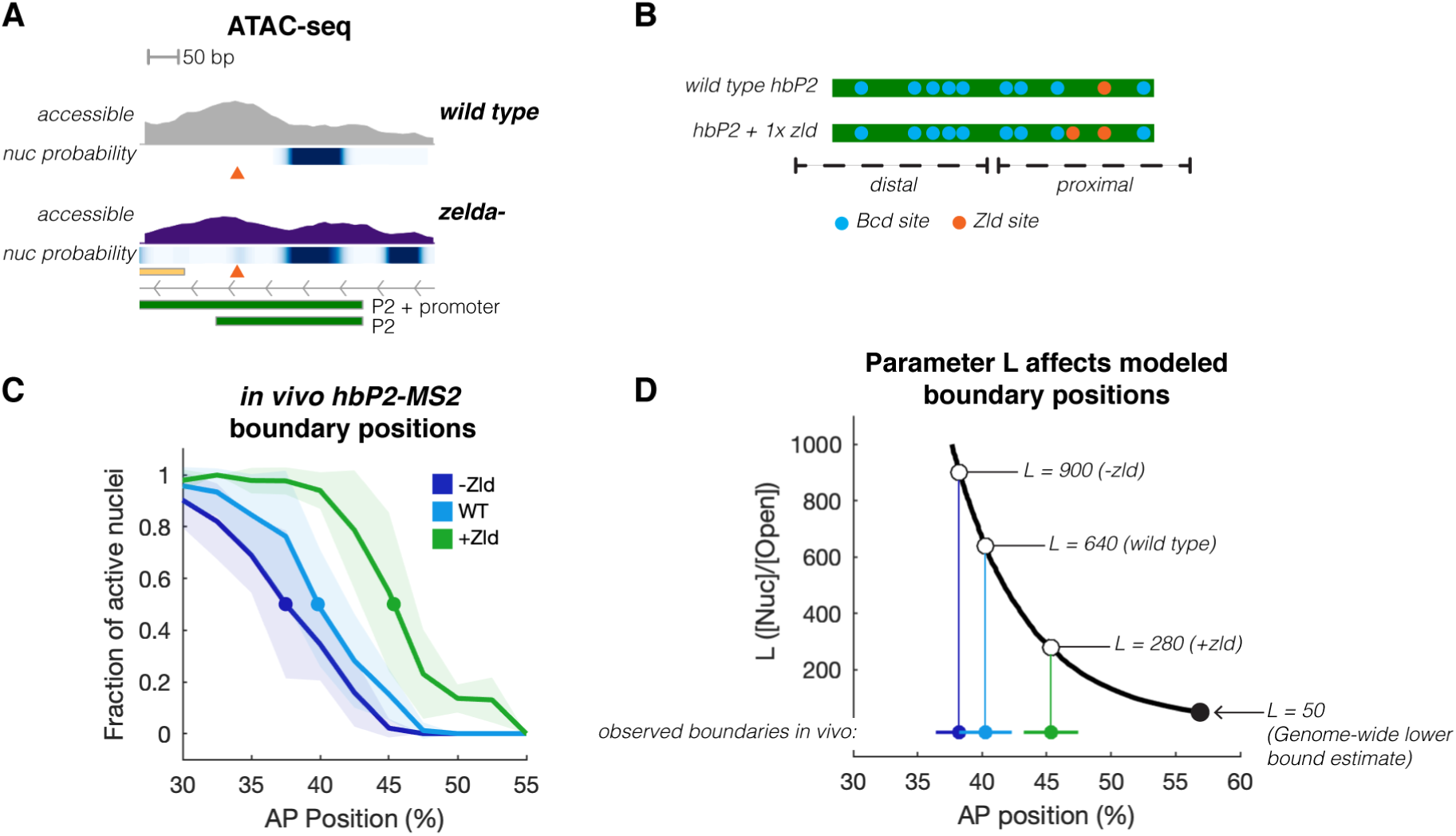
Zelda-dependent changes in reporter activity are approximated by altering modeled nucleosome stability. **A)** Loss of *zelda* has a moderate impact on chromatin organization at *hb* P2. Shown are ATAC measurements of accessibility and modeled nucleosome dyad positions for wild type (top) or *zelda* mutant embryos (bottom). Wild type data is re-plotted from Figure 1B for comparison. Compared with wild type, *zelda* mutants show a modest reduction in chromatin accessibility over *hb* P2 and a slight increase in the occupancy of the proximal nucleosome (orange arrowhead). **B)** Schematic of *hb* P2 minimal elements highlighting the position of the single endogenous Zelda binding site (orange) within the domain of the proximal nucleosome (below), and relative to Bcd binding sites (blue). To experimentally increase the influence of Zelda on this locus, we designed a mutant reporter that includes one additional Zelda site at the indicated position ([*hbP2 + 1x Zld*] *MS2*). **C)** Alteration of Zelda activity at the *hb* P2 reporter shifts the posterior boundary of *hbP2-MS2* expression. Shown is the fraction of active nuclei in 2.5% egg length bins for *hbP2-MS2* in wild type embryos (light blue, n = 19) and *zelda*-RNAi embryos (dark blue, n = 15), as well as for [*hbP2 + 1x Zld*] *MS2* in wild type embryos (green, n = 17). Dots mark the EC50 of each of the fraction of active nuclei profiles determined by Hill equation fits (Supplemental Information). Decreasing Zelda activity with RNAi shifts the posterior boundary of the reporter anteriorly, while increasing Zelda activity at the reporter by adding a Zelda binding site to it shifts the posterior boundary posteriorly. **D)** The effect of experimental modulation of Zelda activity on the measured posterior boundaries can be modeled by changing the nucleosome stability parameter L. Shown are the posterior boundary positions (x axis) that are simulated by the model with specific L values (y axis), as determined by the EC50’s of Hill fits to the simulated fraction of active nuclei (Supplemental Information). The solid black dot indicates the maximal AP position achievable by the model when L is set to the lower bound estimate of 50. Colored dots mark the measured posterior boundaries of of *hbP2-MS2* in wild type (light blue ± standard deviation), *hbP2-MS2* in *zelda*-RNAi (dark blue ± standard deviation) and [*hbP2 + 1x Zld*] MS2 in wild type (green ± standard deviation). *L* = 640, *L* = 900, and *L* = 280 allow for prediction of these measurements respectively.

We applied the two-state model to examine whether alteration of modeled nucleosome stability could result in shifts in the modeled posterior boundary of *hbP2-MS2* expression. In the model, the dimensionless parameter *L* ([Nuc_0TF_]/[Open_0TF_]) represents the equilibrium between the nucleosomal and open states while no TFs are bound to the locus (Mirny, 2010). *L* is termed the “nucleosome stability parameter,” as higher stability nucleosomes will more often occupy DNA, increasing the nucleosome equilibrium occupancy. We estimated a lower bound for *L* as 50-60 based on mitotic ATAC-seq measurements (e.g., Fig 1F, top) of genome-wide fractions of nucleosome associated and open chromatin (Supplemental Information). Our estimate agrees well with prior lower bound estimates of human genome accessibility (range 10-100) based on DNase hypersensitivity measurements (Mirny, 2010). We approximate the upper bound of *L* as 1000, a value previously calculated from nucleosome occupancy measurements (Mirny, 2010). The results of a parameter sweep from *L* = 50 to 1000 reveals that, as *L* increases, the predicted posterior boundary of expression shifts towards the anterior (Fig. 5D). Changing *L* allows the model to simulate the posterior boundaries observed in each of our experimental datasets while holding all other parameters at their best-fit values (*K_O_* = 9 nM, *K_N_* = 2610 nM, and *V_max_* = 1) (Fig. 5D). The model requires a high nucleosome stability value to simulate the effects of decreasing Zld activity (Fig. 5D, dark blue), while it requires a low nucleosome stability value to simulate the effects of increasing Zld activity (Fig. 5D, green). An intermediate nucleosome stability value of *L* = 640 predicts the boundary of *hbP2-MS2* in wild type (Fig. 5D, light blue). Therefore, the range of L seems to define a range of possible posterior boundary positions for a given arrangement of Bcd motifs and nucleosomes. The success of the model in simulating the results of experimentally perturbing Zld activity supports the conclusion that nucleosome stability determines the degree to which transcription depends on Bcd concentration. The model’s versatility underscores the importance of Bcd-nucleosome competition in setting the Bcd concentration threshold for transcription.

### DNA replication delays the onset of transcription at high Bicoid concentrations

While defining *k_on_* according to Bcd-nucleosome competition allows the model to simulate the positioning of the posterior boundary of *hbP2-MS2* expression, the model (Fig. 4D) notably fails to predict accurately the observed timing of transcriptional onset in the anterior (Fig. 2D). The model underestimates both the mean and variance of the onset times at high Bcd concentrations. Regardless of *K_O_, K_N_*, and *L* combination tested, a portion of nuclei simulated in the anterior still initiate transcription prior to the 4.5-minute onset time minimum we observe *in vivo* (Supplemental Information). The discrepancies between the simulations and the *in vivo* data motivated us to dig deeper into the mechanism of concentration-sensitive transcriptional regulation. DNA replication following mitosis 12 could delay transcriptional activation, as transcription in early *Drosophila* embryos requires the completion of DNA replication (Cho et al., 2022). We therefore sought to estimate the impact of DNA replication on the OFF-ON transition of the two-state model to test whether DNA replication contributes to the timing and variance of *hbP2-MS2* onset times.

Active replication origins are thought to be specified at random positions in syncytial embryos (Blumenthal et al., 1974). Therefore, between nuclei A and B from the same embryo, a fixed genomic position (e.g., a reporter) will have variable distances to flanking origins (*ori_1_, ori_2_*) and will therefore complete replication and—by extension—become competent to be transcribed at different times (Fig. 6A, left panel, after Blumenthal et al. 1974). To account for this effect, we designed a new two-state model where switching to the ON state requires both that Bcd bind the reporter and that the promoter completes replication (Fig. 6A, right panel). This replication-dependent model still estimates Bcd binding using the nucleosome-dependent *k_on_*(*[Bcd]*(*x*,*t*)), as described in Figure 4. However, if Bcd evicts nucleosomes prior to the completion of replication at the promoter, the promoter does not transition to ON at that time, and the simulation sets the onset time to the DNA replication time (Fig 6A, right panel). If replication occurs before Bcd evicts the nucleosomes, then Bcd binding instead dictates the promoter’s transcriptional onset time (Fig 6A, right panel). See the Supplemental Information for simulation details.

**Figure 6:**
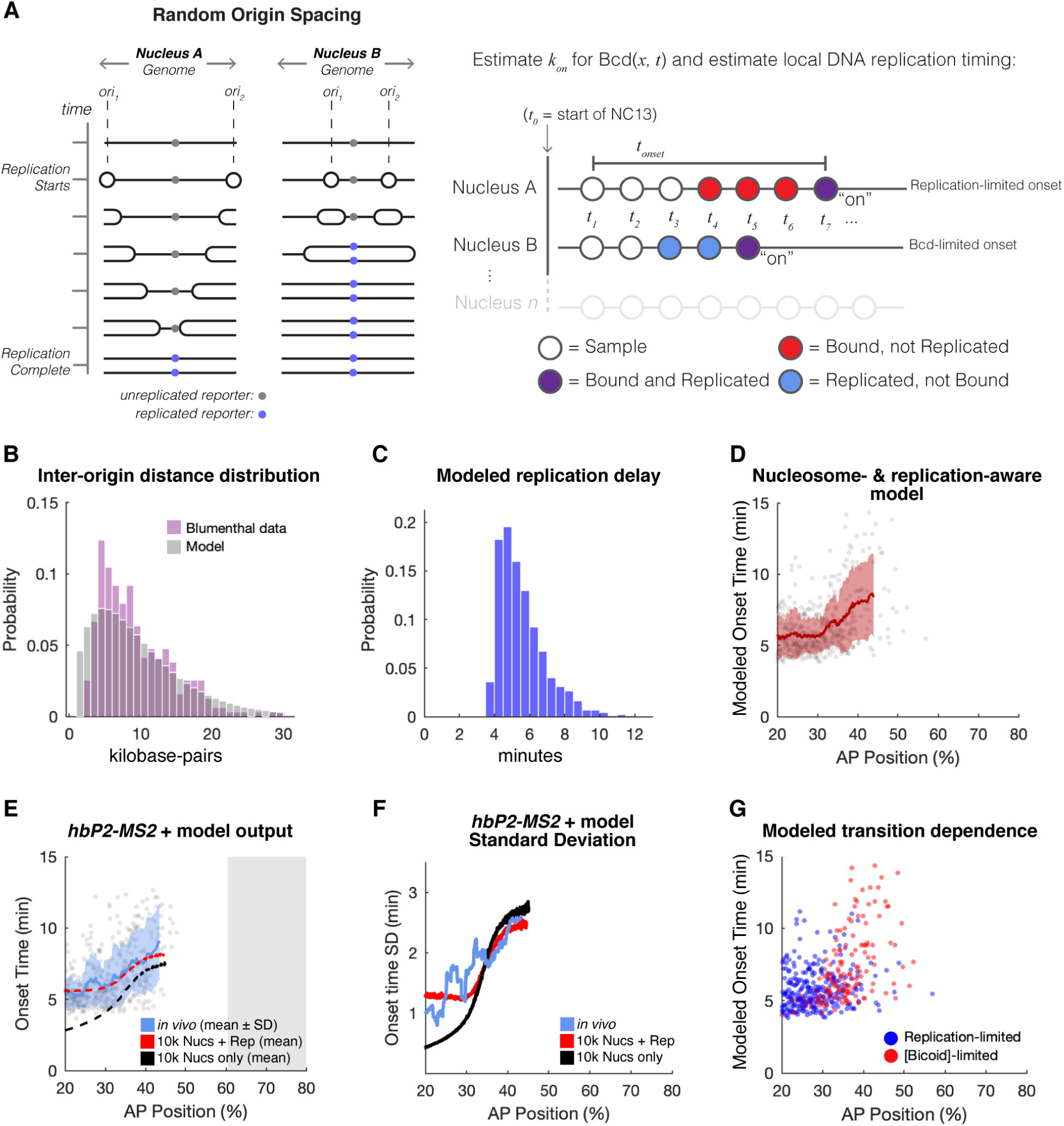
Accounting for the influence of DNA replication on transcription improves the nucleosome-aware model’s predictions. **A)** Cartoon schematic of the revised simulation strategy demonstrated on two example nuclei. As described in the text, the cartoon schematic shows, at left, a single genomic locus in two different nuclei for time points spanning the period of DNA replication, initiated locally at two randomly positioned origins (*ori_1_* and *ori_2_,* after Blumenthal et al 1974). A consequence of random origin spacing is that a fixed genomic locus completes replication at different times between nuclei. Due to the dependency of transcriptional elongation on completion of DNA replication, random origin spacing is expected to impact observed onset times. At right is a revised model schema, which now requires the enhancer both to bind Bcd and to complete replication prior to switching to the ON state. **B)** A measurement of mean origin spacing allows for modeling inter-origin distances. As described in the text, defining the parameters of a Gamma distribution using a 9.7 kb average inter-origin spacing and origin frequency of 2 origins per 9.7 kb allows for estimation of measured inter-origin distances, plotted as a histogram. Measured inter-origin distances were manually measured from Blumenthal et al. 1974 Figure 4 and reproduced here. **C)** Estimation of inter-origin distances allows for prediction of delays in transcriptional onset. The modeled inter-origin distances were converted to replication delays by dividing the distances by 5.3 kb/minute and adding approximately 4 minutes to account for the estimated delay between anaphase and replication onset, plotted as a histogram. **D)** Incorporating replication delays into the nucleosome-aware model alters the onset time distribution predictions. Shown is one simulation of onset times, plotted as a function of AP position. A rolling average of simulated onset times (n = 50) is shown (blue ± standard deviation) across the 95% C.I. of the AP positions of simulated active nuclei. The simulation was performed with *K_O_* = 9 nM, *K_N_*= 2810 nM, and *L* = 600. **E)** A nucleosome- and replication-aware stochastic model accurately predicts the onset time distribution and posterior border of *hbP2-MS2* activity. Shown is a set of sampled *hbP2-MS2* onset times as a function of AP position, with the rolling average of observed onset times (blue ± standard deviation) and the average of 10,000 onset time simulations (red, dotted) using the Nucleosome + Replication model. For comparison, 10,000 simulations with the Nucleosome-only model is shown (black, dotted). Observed and simulated onset time averages are plotted across the 95% C.I. of the AP positions of active nuclei (observed and simulated, respectively). Simulations were performed with *K_O_* = 9 nM, *K_N_* = 2810 nM, and *L* = 600 for both the replication-dependent and independent models. **F)** The nucleosome- and replication-aware stochastic model better accounts for the observed variance in onset time distributions. Plotted are the standard deviations of onset times as a function of AP position for *in vivo* measurements (blue), and the average standard deviations of 10,000 simulations of the Nucleosome + Replication (red) or Nucleosome-only (black) models. **G)** Onset times of *hbP2-MS2* expression are limited either by replication timing or Bcd concentration. Shown is one instance of the simulation (the same as in D), with points indicating the simulated onset times as a function of AP position. The points have been color coded to indicate whether the onset time was limited by either replication time (blue) or Bcd concentration (red).

The model estimates DNA replication timing by drawing from a modeled distribution of inter-origin distances and calculating the time to complete replication. To model replication timing of the reporter, we needed to estimate four parameters: global timing of replication initiation, the speed of DNA polymerase, the relative spacing of active replication origins, and the distribution of origin firing times. We estimated the timing of replication initiation to be 3.75 minutes following anaphase 12 based on prior imaging of PCNA-GFP, which forms foci at sites of active replication whose intensities peak at this time (Blythe & Wieschaus, 2016; McCleland et al., 2009). The speed of DNA Polymerase was previously estimated to be 5.3 kb/min for bidirectional replication from an origin (Blumenthal et al., 1974). Next, while no genomic studies have mapped origin positions specifically in cleavage stage *Drosophila* embryos, the average spacing between origins has been approximated to be 9.7 kb by measuring between replication fork centerpoints observed on electron micrographs. Finally, the statistical distribution of DNA fibers with active replication forks was found to be consistent with synchronous origin firing (Blumenthal et al., 1974). Based on these reported measurements, we estimate that *x* kb between origins, with average spacing of 9.7 kb, takes *x*/5.3 minutes to complete replication following initiation at 3.75 minutes past anaphase 12.

We modeled the likelihood of origin-origin distances by using a gamma distribution with α = 2 origins and β = 9.7 kb/2 origins to estimate the probability that stretches of DNA of length *X* contain a pair of origins:

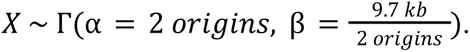

By KS test, we find that the observed origin distribution measurements are indistinguishable from this gamma distribution model at a significance threshold of p = 0.05 (Fig. 6B, Supplemental Information). A gamma distribution describes the waiting times, or in this case distances, before a specified number of events in a Poisson process. The success of the gamma distribution in modeling the measured origin-origin distances supports the assumption of random distribution of origins along chromosomes, with origins activating according to a Poisson process with rate λ = 2 origins/9.7 kb. By dividing our modeled origin-origin distances by 5.3 kb/min and accounting for the four-minute delay before initiation of DNA replication following anaphase 12, we calculated a distribution of possible times at which the reporter’s promoter completes replication (Fig. 6C). This replication time distribution allows the two-state model of promoter activity to account for transcription’s requirement for DNA replication.

We find that the DNA replication, nucleosome-dependent model of transcriptional activation successfully recapitulates the measured *hbP2-MS2* onset time distribution (Fig. 6D & E). The OFF-ON transition’s requirement for the completion of DNA replication at the promoter introduces additional variance into the simulated onset times at high Bcd concentrations that better reflects the variance observed in our *hbP2-MS2* data (Fig 6F). Furthermore, the requirement for DNA replication prevents anterior onset time predictions substantially prior to five minutes into NC13. The new model structure also accurately predicts the posterior boundary of MS2 expression; the model’s best-fit parameter set contains Bcd-nucleosome competition parameters that place the posterior boundary of simulated expression at 45% EL (*K_O_* = 9 nM, *K_N_* = 2810 nM, and *L* = 600) (Fig. 6D & E). The model suggests DNA replication delays transcriptional onset at the high Bcd concentrations where Bcd readily outcompetes nucleosomes, while at lower Bcd concentrations the requirement that Bcd outcompetes nucleosomes dictates the timing of transcriptional onset (Fig. 6G).

### Genome replication and Bicoid binding regulate RNA polymerase II pause-release

We make two assumptions in the above onset time simulation about the relationship between TF binding, RNA Pol II, and DNA replication. We assume that, following mitotic exit, Bcd has the ability to compete with nucleosomes and bind to DNA independently of the completion of DNA replication. An alternative possibility, however, is that DNA replication could itself lower the nucleosome barrier for TF binding, as passage of the replication fork disrupts nucleosomes and transiently reduces the nucleosome content of DNA by half (Weintraub, 1974). We also assume that transcription becomes immediately detectable following the completion of both Bcd binding and DNA replication, and that therefore the recruitment of RNA Pol II to the promoter likely occurs independently of these processes (Fig 6A). The transition of RNA Pol II from transcriptional initiation to elongation, akin to a pause-release, in contrast may require Bcd binding and the completion of DNA replication. We experimentally tested the following hypotheses that stem from our assumptions: 1) Bcd binding to *hb* P2 occurs independently of DNA replication, 2) Pol II recruitment to *hb* occurs independently of DNA replication, and 3) Pol II recruitment to *hb* occurs independently of Bcd binding.

To determine how DNA replication impacts Bcd-dependent transcription, we measured Bcd and Pol II occupancy in control and DNA replication-inhibited embryos. To collect DNA replication-inhibited embryos for ChIP-seq, we developed a protocol for treating embryos in bulk with hydroxyurea (HU) to block their synthesis of dNTPs (Fig S2). ChIP-seq measurements do not reveal any differences in Bcd binding at *hb* P2 between control and HU-treated embryos (Fig. 7A, blue). Genome-wide Bcd binding also does not change substantially upon inhibition of DNA replication (Fig. 7B). Therefore, Bcd binds to DNA independently of DNA replication state, and its ability to outcompete nucleosomes likely does not depend on the disruption of chromatin by replication forks. In contrast, RNA Pol II coverage across gene bodies is strongly affected when DNA replication is inhibited. At the *hb* locus, RNA Pol II is markedly reduced over the gene body, yet retains a small peak at the P2 transcriptional start site, consistent with a replication-sensitive block to the transition between initiation and elongation (Fig. 7A, red). Genes expressed prior to large-scale zygotic genome activation (“pre-MBT” genes (K. Chen et al., 2013)) also exhibit a higher ratio of initiated vs. elongating Pol II upon replication inhibition (Fig. 7C-C’, Fig. S3). These measurements confirm the central assumptions of our model, and indicate that the completion of DNA replication precedes the transition from Pol II initiation to elongation.

**Figure 7:**
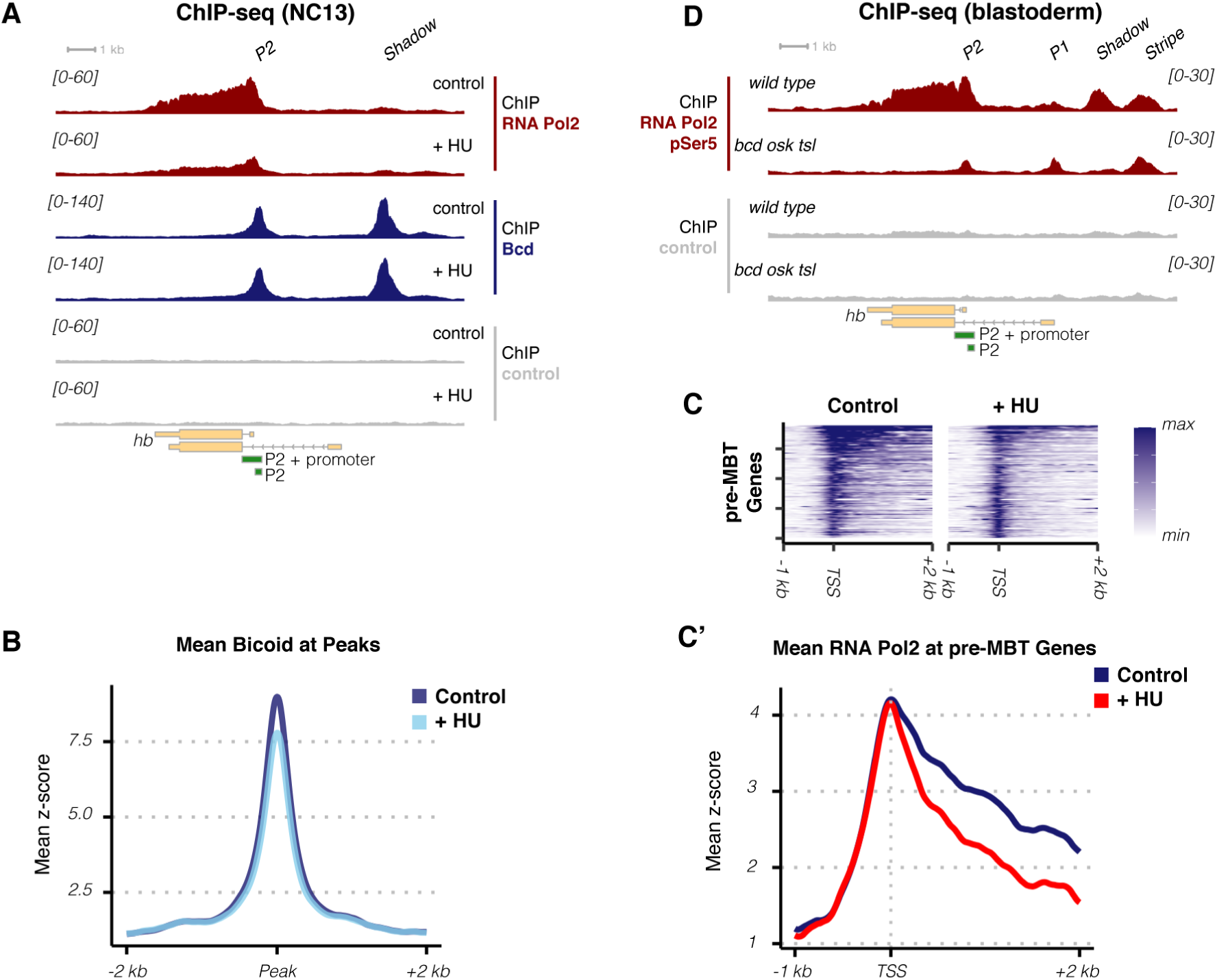
Initiated RNA Pol II waits for the completion of replication and Bcd binding to enter into productive elongation of *hb*. **A)** Inhibition of DNA replication does not impact Bcd binding at the *hb* locus, but reduces Pol II occupancy over the gene body. Shown is ChIP-seq data over the *hb* locus for RNA Pol II (dark red), Bcd (blue), and control (gray) comparing occupancy at NC13 in control (untreated) and HU-treated embryos. Data is the average of two independent biological replicates, normalized to read depth (counts per million reads, CPM). The y-axis range for each plot is shown at left (brackets). **B)** Inhibition of DNA replication does not significantly affect genome-wide Bcd binding. Bcd ChIP-seq data for control (dark blue) and HU-treated (light blue) embryos was standardized and averaged over 1,026 peak regions. The plot shows the average standardized Bcd binding over these peaks. **C)** Inhibition of DNA replication in NC12-13 primarily affects the entry of RNA Pol II into productive elongation. Upper panels show heatmap representation of standardized RNA Pol II ChIP-seq signal over a set of genes transcribed before large-scale ZGA (pre-MBT genes, Chen et al, 2013). Compared with the control, HU treatment results in loss of RNA Pol II within gene bodies, but retention of a peak near the TSS. The lower panel (C’) is the average of the heatmap representation, with HU-treatment (red) demonstrating reduced Pol II in the gene body but retention of signal over the TSS compared with control (dark blue). **D)** RNA Pol II at *hb* is maintained in a paused state in the absence of Bcd. ChIP-seq for initiated RNA Pol II (pSer5) or control was performed on wild type or *bcd osk tsl* blastoderm stage embryos. In the absence of Bcd, RNA Pol II forms a distinct peak at the *hb* P2 TSS. Data is an average of three independent biological replicates, normalized to read depth (counts per million reads, CPM). The y-axis range for each plot is shown at right (brackets)..

**Supplemental Figure S2, related to Figure 7:**
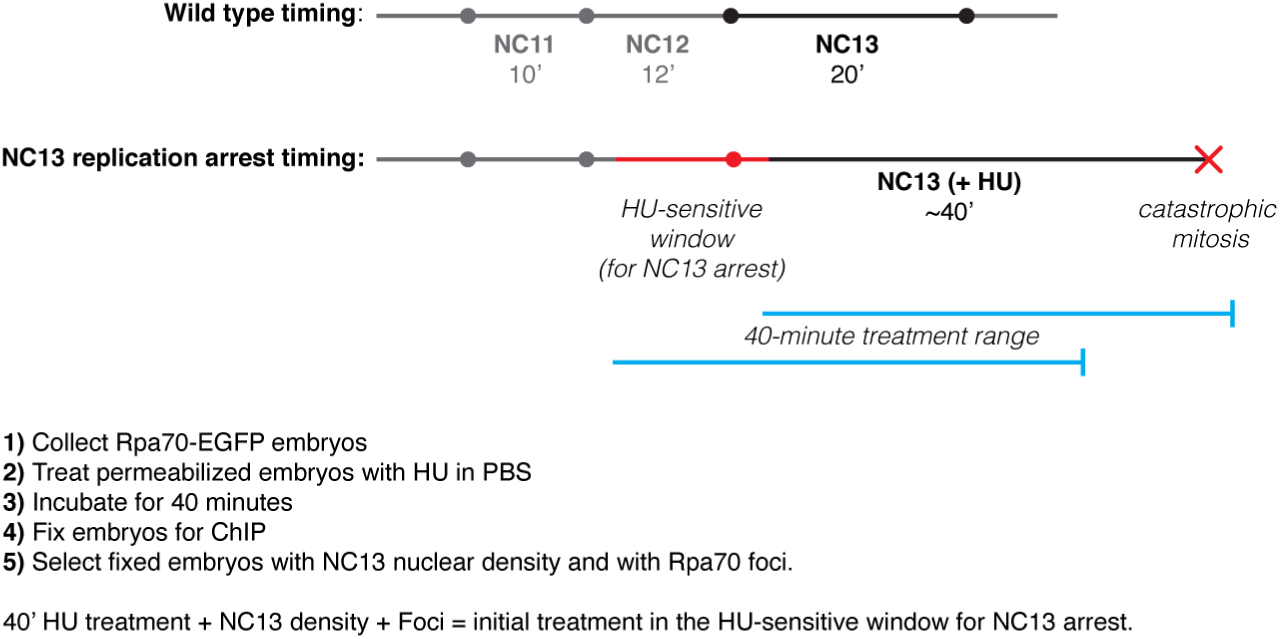
Schematic for HU treatment and collection of NC13 replication arrested embryos. Based on live confocal imaging of the response of embryos to HU treatment (Blythe & Wieschaus, 2015b) exposure of embryos to HU after initiation of replication in NC12 results in replication inhibition during the following cell cycle (NC13). Replication inhibited embryos form foci of RpA-70 and remain in an interphase-like state for nearly 40 minutes before undergoing a catastrophic mitosis with un-replicated DNA. For a bulk collection containing unsynchronized embryos, we reasoned that treating embryos with HU for 40 minutes prior to fixation for ChIP would allow us to select embryos with NC13 nuclear density, and by evaluating the presence of RpA-70 foci, to select the embryos undergoing replication inhibition.

**Supplemental Figure S3, related to Figure 7:**
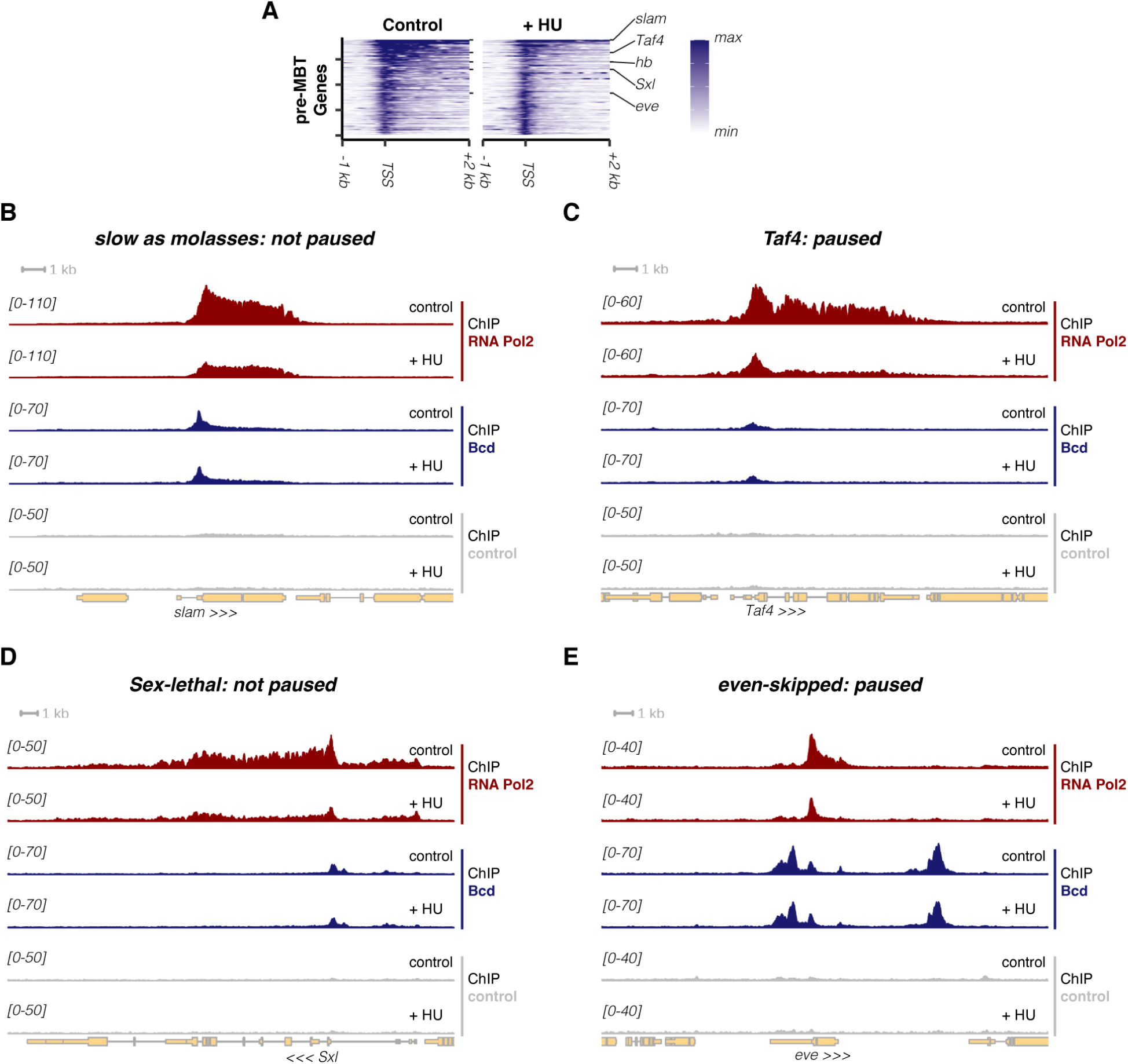
Further examples of Pol II and Bcd binding to pre-MBT transcribed genes following HU treatment. **A)** The heatmap from Figure 7C is reproduced, with the relative locations shown for the five loci detailed at high resolution (Figure 7A, Figure S2B-E). **B)** The *slow as molasses (slam)* locus is detailed. Genes like *slam* and *Sex-lethal* (Fig S2D) do not undergo transcriptional pausing at pre-MBT stages (Chen et al, 2013). Following replication inhibition with HU, RNA Pol II is uniformly reduced over the *slam* coding region. In panels B-E, RNA Pol II (red), Bicoid (blue) and negative control (gray) ChIP-seq coverage is shown over the indicated genomic locus for control and HU-treated embryos. The data are normalized to counts per million total mapped reads, and the displayed y-axis range is indicated in the brackets at left. The directionality of the transcription unit of interest is indicated by carats (<<< or >>>) at the bottom of each plot. **C)** The *Taf4* locus is detailed. Genes like *Taf4* and *even-skipped* (Fig S2E) are subject to transcriptional pausing at the MBT. Following replication inhibition with HU, RNA Pol II is depleted from the *Taf4* gene body, and shows an accumulation at the transcriptional start site. **D)** The *Sex-lethal (Sxl)* locus is detailed. Similar to *slam*, *Sxl* demonstrates uniform reduction of RNA Pol II over the gene body following replication inhibition with HU. **E)** The *even-skipped (eve)* locus is detailed. Similar to *Taf4*, *eve* demonstrates retention of RNA Pol II over the transcription start site following replication inhibition with HU.

**Supplemental Figure S4, related to Figure 7:**
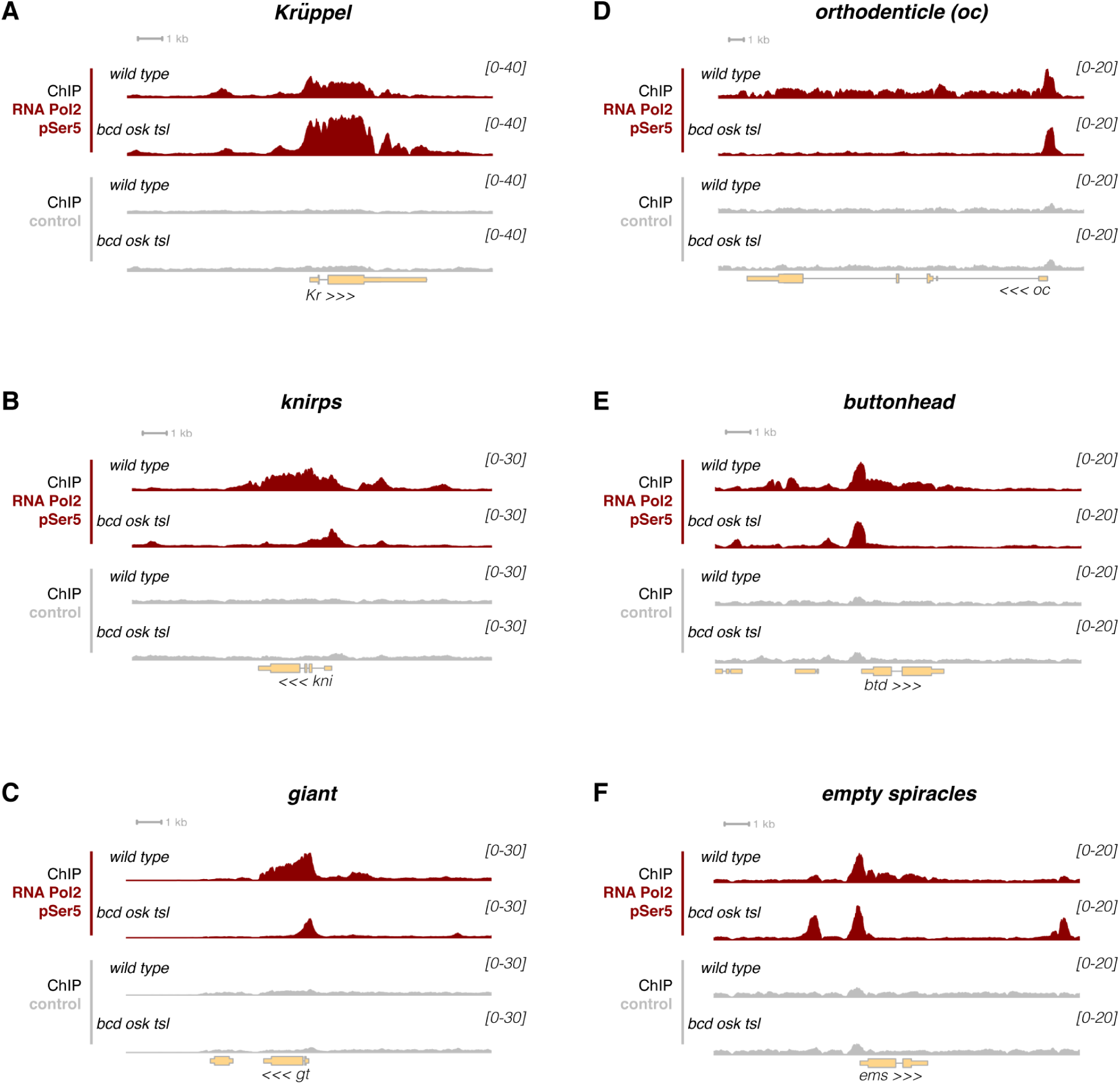
Further examples of Pol II binding to Bcd targets in wild type and *bcd osk tsl* mutant embryos. **A)** The *Kruppel* locus is detailed. *Kruppel* is uniformly expressed in *bcd osk tsl* mutant embryos (Petkova et al., 2019). The RNA Pol II ChIP-seq captures this quantitative effect, showing increased RNA Pol II binding to the *Kruppel* locus in *bcd osk tsl* mutant embryos. For all panels in this figure, RNA Pol II (red) and negative control (gray) ChIP-seq coverage is shown over the indicated genomic locus for wild type and *bcd osk tsl* mutant embryos. The data are normalized to counts per million total mapped reads, and the displayed y-axis range is indicated in the brackets at right. The directionality of the transcription unit of interest is indicated by carats (<<< or >>>) at the bottom of each plot. **B)** The *knirps* locus is detailed. In *bcd osk tsl* mutant embryos, RNA Pol II accumulates at the transcription start sites of the *knirps* locus. **C)** The *giant* locus is detailed. In *bcd osk tsl* mutant embryos, RNA Pol II accumulates at the transcription start site of the *giant* locus. **D)** The *ocelliless/orthodenticle* locus is detailed. In *bcd osk tsl* mutant embryos, RNA Pol II accumulates at the transcription start site of the *orthodenticle* locus. **E)** The *buttonhead* locus is detailed. In *bcd osk tsl* mutant embryos, RNA Pol II accumulates at the transcription start site of the *buttonhead* locus. **F)** The *empty spiracles* locus is detailed. In *bcd osk tsl* mutant embryos, RNA Pol II accumulates at the transcription start site of the *empty spiracles* locus.

We next evaluated the independence of Pol II initiation from Bcd binding by performing Pol II ChIP-seq in wild type embryos and embryos triply mutant for the maternal factors *bicoid* (*bcd*), *oskar* (*osk*), and *torso-like* (*tsl*). We chose to use *bcd osk tsl* rather than *bcd^E1^* embryos, as the triple mutant context eliminates AP and terminal maternal factors that contribute to the regulation of anterior and posterior *hb* stripe enhancers, reducing confounding Pol II activity at *hb*. In triple mutant embryos, we find that while Pol II is lost from the *hb* gene body, Pol II remains associated with the P2 promoter (Fig. 7D). Nearly every Bcd target we assess also maintains Pol II at its promoter in the triple mutant condition (Fig. S4). The preservation of Pol II at promoters indicates transcriptional initiation at *hb* is independent of Bcd activity. Bcd likely primarily activates transcription by increasing the probability of Pol II elongation. The regulation of Pol II pause-release likely underlies how Bcd can rapidly initiate transcription within the short cell cycles of early development.

## DISCUSSION

The molecular mechanisms that determine the concentration thresholds of Bcd target genes have been pursued since the discovery of Bcd in 1988 (H. Chen et al., 2012; Driever et al., 1989; Driever & Nüsslein-Volhard, 1988a, 1988b; Hannon et al., 2017; Ochoa-Espinosa et al., 2005; Struhl et al., 1989). Here, we have illustrated that the pattern of Bcd binding sites and nucleosomes at an enhancer uniquely determines a Bcd concentration threshold for transcription, and have provided a generalizable framework for understanding Bcd concentration-sensitivity. With physically-reasonable parameter values, our computational model of promoter activity accurately simulates how the timing of *hbP2-MS2* transcriptional onset changes across Bcd concentrations. Our theoretical and experimental work asserts Bcd-nucleosome competition sets a concentration threshold for transcription, while DNA replication delays transcriptional onset at high Bcd concentrations where Bcd “wins” its competition with nucleosomes. Experimentally testing the model’s predictions through live-imaging *hbP2-MS2* both validates the model and reveals nucleosome stability may determine the degree to which the posterior boundary of *hb* transcription depends on Bcd concentration. ChIP-seq experiments also support the model’s structure, demonstrating that establishment of active transcriptional elongation at *hb* and elsewhere depends both on Bcd binding and completion of DNA replication. We conclude that Pol II elongation on a Bcd target gene following mitosis requires both that Bcd outcompetes nucleosomes for occupancy at an enhancer and that an associated promoter completes DNA replication. This *hbP2* case-study study provides a molecular explanation for how the *in vivo* environment of early development shapes the formation of concentration-sensitive patterning gene expression domains.

TF-enhancer interactions must be understood in the context of chromatin and the cell cycle. Nucleosomes act as barriers to TF binding to genomic loci and rapid cell divisions limit transcriptional outputs early in the development of many organisms (Blythe & Wieschaus, 2015a; Kaplan et al., 2009; Kornberg, 1974; Tadros & Lipshitz, 2009). This work fills in gaps in understanding highlighted by prior models that describe in the abstract how the maternal factors Bcd and Zld regulate a promoter’s OFF-ON transition (Dufourt et al., 2018; Eck et al., 2020; Fernandes et al., 2022). A prior model found that Zld shortens the duration of inactive promoter states preceding transcription of a reporter for a DV patterning enhancer (Dufourt et al., 2018). Zld similarly influences the OFF-ON transition in our model, pointing to the conclusion that the previously characterized inactive promoter states correspond to high degrees of nucleosome occupancy at the enhancer. At both DV and AP target genes, Zld likely speeds the transition between inactive states by acting as a pioneer factor and by reducing nucleosome stability at enhancers. In order to account for the delays preceding transcription of a *hb* P2 + P2 promoter reporter, another model incorporated irreversible transitions through inactive promoter states driven by Bcd and Zld (Eck et al., 2020). Here, we specify that the molecular mechanisms that account for the irreversible OFF-ON transition are Bcd-nucleosome competition and the dependence of Pol II elongation on the completion of DNA replication. Recently, a model based on synthetic enhancer measurements suggested that Zld speeds a promoter’s OFF-ON transition by increasing the rate of Bcd binding (Fernandes et al., 2022). This finding fits within the theoretical framework we present, as Zld’s impact on nucleosome stability allows Bcd to more easily outcompete nucleosomes and bind its sites. Importantly, our work specifies that at high TF activity levels, the timing of DNA replication at a locus sets an upper limit for the rate of transcriptional activation.

This work fits within expectations based on prior observations of Bcd dynamics within the nucleus and the formation of clusters at presumed binding sites (Kawasaki & Fukaya, 2023; Mir et al., 2017, 2018; Munshi et al., 2024). A broad range of TFs have been observed to form high concentration assemblies or clusters within the nucleus (Boijja et al., 2018; Dufourt et al., 2018; Kawasaki & Fukaya, 2023; Mir et al., 2017, 2018; Munshi et al., 2024; Wollman et al., 2017). In the case of Bcd, while clusters contain only a small fraction of total protein, they show strong spatial and temporal correlation with target gene transcription as measured with an MS2 reporter (Mir et al., 2018; Munshi et al., 2024). How the frequencies of cluster formation and dispersal impact the dynamics of Pol II activity at a promoter remains an open question. A recent study estimated the number of Bcd molecules within clusters coinciding with the *hb* P2 expression domain to be greater than or equal to the number of expected binding events at *hb* P2 (≥9) (Munshi et al., 2024). In light of our work, we propose that Bcd clustering allows for rapid delivery of the high local concentrations of Bcd necessary to outcompete nucleosomes for occupancy at the nine Bcd sites in *hb* P2. Therefore, the frequency of cluster formation provides an upper limit for the rate at which Bcd molecules have the opportunity to outcompete the nucleosomes at a locus. As our model’s OFF-ON transition rate depends on Bcd’s ability to outcompete nucleosomes, it provides a framework for translating TF cluster dynamics to Pol II behavior at a promoter. We propose periodic TF cluster formation regulates the transcriptional process by incrementally reorganizing the chromatin at enhancers.

Finally, this work raises the possibility that the input-output relationship between any TF and a target gene could be modeled accurately, given nucleosome positions measured both in the presence and absence of key regulators. In the case of Bcd, we expect that this model will, with elaboration, explain the mechanisms producing the concentration thresholds of additional target genes. Due to its apparent lack of additional regulatory inputs, *hb* P2 represents a simplified target of investigation. Other physiologically important Bcd targets receive competitive input from repressors and undergo more extensive chromatin reorganization by pioneers such as Zld (H. Chen et al., 2012; Hannon et al., 2017). With additional work, an allosteric modeling framework could incorporate the activity of repressors and pioneers that—either directly or indirectly—modulate an activator’s ability to access its sites. Further, we conceptualize nucleosome occupancy as a means to create suboptimal, effectively low affinity TF binding conditions. Our model therefore supports previous observations of the importance of low affinity TF binding motifs for spatially-restricted enhancer activity in response to graded and uniform TF inputs (Crocker et al., 2015; Farley et al., 2015; Ramos & Barolo, 2013). We hypothesize that, across organisms and developmental contexts, nucleosomes play key gene regulatory roles by creating conditions that require high TF concentrations for TF-DNA binding.

## MATERIALS AND METHODS

### EXPERIMENTAL MODEL DETAILS

#### Drosophila stocks

This study used the following *Drosophila melanogaster* (NCBI Taxon 7227) stocks.

*w^1118^* (source: BestGene Inc., Chino Hills, CA).

*y {w^+^= His2Av-miRFP670-T2A-HO1}ZH2A w* (this study)

*w; {w^+^= RpA-70-EGFP}attP2* (Blythe & Wieschaus, 2015b) *bcd^E2^ osk^166^ tsl^4^/TM3*

*yw; P*{w=*his2AV-RFP*}2*; P{w=MCP-GFP}4F* (kind gift of H. Garcia) (Eck et al., 2020)

*y w; {w^+^=nos>MCP-mCherry}VK22* (kind gift of H. Garcia)

*cn bw Kr^1^/SM6 bw* (Bloomington *Drosophila* Stock Center (BDSC) #3494)

*w; UASp>zldRNAi (III)* (kind gift of C. Rushlow) (Sun et al., 2015)

*w;* P{*w=alpha-tubulin67c-GAL4 VP16*}67 (Princeton Stock Collection)

*w; bcd^E1^/TM3* (Princeton Stock Collection)

*w* P{*w=His2Av-RFP*}1; *bcd^EGFP^* (kind gift of P. Onal and S. Small)

All fly stocks were maintained on an enriched high-agar cornmeal media as reported previously (Cline, 1978).

### Transgenic lines

A *hbP2-MS2* reporter was made using the *hb* P2 minimal enhancer, 245 bp (chr3R:8,694,654-8,694,898, dm6 reference assembly) from the 263 bp *hb* regulatory element identified previously (Struhl et al., 1989). The *hb* P2 sequence was cloned upstream of the Drosophila Synthetic Core Promoter (Pfeiffer et al., 2008) which drives expression of 24 MS2(v5) stem loop sequences (Yoon et al., 2016) embedded in the intron of *yellow* and followed by the *Gal4* sequence in an attB transgenic vector. The reporter was integrated into the attP40 landing site on the second chromosome using PhiC31 integrase-mediated transgenesis. A *[hbP2 + 1xZld]-MS2* reporter was made by introducing an optimal *zld* motif to base pair positions 178-185 within *hb* P2, a location that does not disrupt any of the TF binding sites in *hb* P2 identified with PWM matches. Besides the *zld* motif alteration to hbP2, the *[hbP2 + 1xZld]-MS2* line was made through an identical cloning and integration process to that performed to make *hbP2-MS2*.

A transgenic line that produces embryos that express Bcd uniformly was achieved as follows. First, a *w;* **α***Tub67C-EGFP-Bcd/^FRT^bcd 3’UTR 3xP3>dsRed/^FRT^sqh 3’UTR/TM3 (III)* transgenic line was made by inserting a previously published uniform Bcd construct (Hannon et al., 2017) into the VK33 landing site on the third chromosome using PhiC31 integrase-mediated transgenesis. The **α***Tub67C-EGFP-Bcd/^FRT^bcd 3’UTR 3xP3>dsRed/^FRT^sqh 3’UTR* transgene was then recombined with the *bcd^E1^* allele to produce a **α***Tub67C-EGFP-Bcd/^FRT^bcd 3’UTR 3xP3>dsRed/^FRT^sqh 3’UTR, bcd^E1^* recombinant. Uniform Bcd expression was achieved as described previously (Hannon et al., 2017) by excising the FRT-flanked *bcd* 3’UTR with transient induction of Flp expression and scoring for loss of the *3xP3>dsRed* marker. This yields **α***Tub67C-EGFP-Bcd sqh 3’UTR, bcd^E1^*, which expresses Bcd uniformly is henceforth referred to as **α***Tub67C-uEGFP-Bcd, bcd^E1^*.

### Datasets

The ChIP- and ATAC-seq data for wild-type versus uniform Bicoid, including predicted nucleosome dyad positions were previously reported (Hannon et al., 2017) (GSE86966). ATAC-seq data from different stages of the NC13 cell cycle were previously reported (Blythe & Wieschaus, 2016) (GSE83851). The processed Bcd ChIP-Nexus data demonstrating positive and negative sequence contributions to Bcd binding (Brennan et al., 2023) was retrieved from https://zenodo.org/record/8075860 and plotted over the *hb* genomic locus.

Data from the replication inhibition ChIP-seq experiment and Pol II binding in *bcd osk tsl* will be submitted to GEO-SRA.

## METHOD DETAILS

### Live-imaging

#### MS2 reporters

Embryos for *hbP2-MS2* and *[hbP2 + 1xZld]-MS2* imaging: Embryos were collected from a cross of *hbP2-MS2* or *[hbP2 + 1xZld]-MS2* males to *yw/w; his2AV-RFP/+; MCP-GFP/+* females. Embryos for *hbP2-MS2* in uniform Bcd imaging: Embryos were collected from a cross of *hbP2-MS2* males to *his-miRFP670/+; MCP-mCh/+;* **α***Tub67C-uGFP-Bcd, bcd^E1^/bcd^E1^* females. Embryos for *hbP2-MS2* in zld^RNAi^ imaging: Embryos were collected from a cross of *hbP2-MS2* males to *his2AV-RFP/+; mat-tub-GAL4 67/+; UAS-shRNA-zld/+* females. The imaging of MS2 transcription in all genotypes was performed as follows.

Pre-NC13 embryos were dechorionated in 4% bleach (50% dilution of concentrated Clorox) for 1-2 min. Dechorionated embryos were mounted in Halocarbon 27 oil (Sigma) on a gas-permeable membrane (BioFoil, Heraeus, New Jersey, USA) and overlaid with a glass coverslip. Embryos were imaged on a Leica SP8 WLL Confocal Microscope using a 63x 1.3NA glycerol objective, 1.28x zoom, and 700 Hz scan rate at a 512×256 pixel resolution. When imaging MS2 in graded Bcd embryos, a 590 nm laser (measured as 15 μW through the 10x objective) and a 488 nm laser (measured as 42 μW through the 10x objective) were used to excite RFP and GFP respectively. RFP and GFP emissions were collected at [602, 650] nm (detector set to BrightR) and [498, 531] nm (detector set to Photon Counting) respectively. When imaging MS2 in uniform GFP-Bcd embryos, a 488 nm laser (measured as 29 μW through the 10x objective) was used to excite GFP-Bcd, a 587 nm laser (measured as 18 μW through the 10x objective) was used to excite MCP-mCh, and a 670 nm laser (measured as 56 μW through the 10x objective) was used to excite His2Av-miRFP670. GFP-Bcd emission was collected at [498, 535] nm (detector set to Photon Counting), MCP-mCh emission was collected at [597, 629] nm (detector set to Photon Counting), and his-miRFP670 emission was collected at [680, 720] nm (detector set to BrightR). As the maximum laser outputs of confocal microscopes can fluctuate day-to-day, the % outputs of the lasers were adjusted prior to each imaging session to ensure that embryos were imaged with equivalent laser powers between imaging sessions. For all genotypes, per embryo, a 10 μm z-stack was imaged at 10 s/frame and with a 0.5 μm z-step from pre-NC13 to at least 15 minutes into NC14. Immediately following live-imaging of MS2 in graded Bcd embryos, an overview image of the embryo was taken by performing a tile scan using 590 nm excitation at 700 Hz and [602, 650] nm detection with a top z-plane intersecting the center of nuclei and a bottom z-plane at the embryo’s midsagittal plane. Tiles were merged into a single image on the Leica software. Immediately following live-imaging of MS2 in uniform GFP-Bcd, an overview image was taken by following an equivalent procedure, with the exception of using a 488 nm laser and 200 Hz scan speed for exciting GFP and detecting emission at [498, 535] nm.

#### GFP fusion proteins

For live-imaging *His2AV-RFP; bcd^EGFP^* embryos, pre-NC13 embryos were dechorionated and mounted for imaging through the aforementioned MS2 imaging method. Embryos were imaged on a Leica SP8 WLL Confocal Microscope using a 63x 1.3NA glycerol objective, 1.28x zoom, and 700 Hz scan rate at 512×256 pixels. A 590 nm laser (measured as 19 μW through the 10x objective) and a 488 nm laser (measured as 28 μW through the 10x objective) were used to excite RFP and GFP respectively. RFP and GFP emissions were collected at [602, 650] nm (detector set to BrightR) and [499, 531] nm (detector set to Photon Counting) respectively. Per embryo, a 10-μm z-stack was imaged at 10 seconds/frame and with a 0.5 μm z-step from pre-NC13 to at least 15 minutes into NC14.

### Hybridization chain reaction

#### Collection

To allow for measurement of *hbP2-MS2* in *Kr* mutant embryos, the *hbP2-MS2* reporter chromosome was recombined with *cn bw Kr^1^* to generate a *hbP2-MS2 Kr^1^/SM6* line. Embryos were collected from a cross of *hbP2-MS2 Kr^1^/SM6* females to *hbP2-MS2 Kr^1^/SM6* males after a laying period of 4 hr on an apple juice-agarose plate. The mutant embryo genotype produced was *hbP2-MS2 Kr^1^/hbP2-MS2 Kr^1^* and the control embryo genotype was *hbP2-MS2 Kr^1^/SM6*. For measurement of *hbP2-MS2* in embryos uniformly expressing Bcd, embryos were collected from a cross of **α***Tub67C-uEGFP-Bcd, bcd^E1^*/*bcd^E1^* females to *hbP2-MS2* males after a laying period of 4 hr on an apple juice-agarose plate. Control embryos were similarly collected for the uniform Bcd experiment from a cross of *white* females to *hbP2-MS2* males.

#### Fixation

Embryos were dechorionated for 1 minute in 4% bleach and then added to a combination of 1 mL 20% formaldehyde, 4 mL 1x Phosphate Buffered Saline (PBS), and 5 mL heptane. After the embryos were shaken for 20 min, the aqueous layer was removed, methanol was added, and the embryos were vortexed for 30 seconds to remove their vitelline membranes. The embryos were then washed 3x in methanol and stored in methanol at -20 C for at least overnight before undergoing the staining protocol. For the *Kr* mutant experiment, mutant and control embryos were fixed together in the same tube, as they were produced by the same cross and can be distinguished by the presence of Kr mRNA. For the uniform Bcd experiment, mutant and control embryos were fixed separately.

#### Probes and amplifiers

The following RNA probes were synthesized by Molecular Instruments: a probe that targets Gal4 mRNA and is compatible with B1 amplifier sets and a probe that targets Kr mRNA and is compatible with B3 amplifier sets. B1-488 nm, B1-647 nm, and B3-546 nm RNA hairpin-fluorophore conjugates were synthesized by Molecular Instruments and used as amplifiers.

#### Staining

The Hybridization Chain Reaction (HCR) In Situ Protocol V.1 (Bruce et al., 2021) was performed on fixed embryos with the following modifications. For the embryo permeabilization, embryos were permeabilized for 30 minutes at room temperature rocking in a detergent solution (0.1% Triton X-100, 0.05% Igepal CA-630, 500 μg/mL sodium deoxycholate, 500 μg/mLsaponin, 2 mg/mL BSA Fraction V). In our hands, use of SDS for embryo permeabilization as recommended in the cited protocol resulted in undesirable expansion of the embryos and difficulty in downstream handling. We substituted the above detergent mix without loss of signal intensity as determined in a pairwise comparison. For the probe incubation, the embryos were incubated for 17 hr at 37°C in a solution of probe hybridization buffer, 0.8 pmol of Gal4-B1 probe, and 0.8 pmol of Kr-B3 probe (Kr mutant experiment), or for 17 hr at 37°C in a solution of probe hybridization buffer and 0.8 pmol of Gal4-B1 probe (uniform Bcd experiment). At the amplification step, an amplification reaction was performed by incubating the embryos in 0.06 μM of each of the B1-488 nm and B3-546 nm hairpin sets for 24 hr at room temperature (Kr mutant experiment) or in 0.06 μM of the B1-647 nm hairpin set for 18 hr at room temperature (uniform Bcd experiment). Following washes according to Bruce et al., 2021, embryos were stained with 1 ug/mL DAPI (4’,6-diamidino-2-phenylindole; Invitrogen REF: D1306) in 50% glycerol (in 1xPBS) rocking for 1 hr at room temperature. The DAPI solution was replaced with 50% glycerol (in 1xPBS) and the embryos stored at 4°C prior to mounting.

#### Imaging

Embryos were mounted in Prolong Gold Antifade Mountant (Invitrogen P36934) and overlaid with a glass coverslip. Images were collected on a Leica SP8 WLL Confocal Microscope using a 20x 0.75 NA glycerol objective and 400 Hz scan rate at 512×256 pixels and 12-bit resolution. Images were collected at 0.85x zoom (Kr mutant experiment) or 1x zoom (uniform Bcd experiment). A z-stack that spanned from the top of the embryo to the embryo’s midsagittal plane was imaged with a 1.04 μm z-step size. For the Kr mutant experiment, mutant and control embryos were imaged on the same microscope slide and distinguished by the presence of Kr mRNA, as the *Kr^1^*mutant is RNA-null. Gal4 staining was excited with a 488 nm laser and detected at [495, 516] nM (Photon Counting), and Kr staining was excited with a 457 nm laser and detected at [559, 573] nM (Photon Counting). Both channels were collected with a line accumulation of 3. For the uniform Bcd experiment, mutant and control embryos were mounted and imaged on different microscope slides. Gal4 staining was excited with a 650 nM laser (measured as 48 μW through the 10x objective for both mutant and control) and detected at [672, 694] nM (Photon Counting) with a line accumulation of 1. In both experiments, Dapi was excited with a 405 nm laser.

### Chromatin immunoprecipitation

#### Hydroxyurea treatment

Embryos were collected from *RpA70-EGFP* parents on standard apple juice-agar plates after a 2.5-hr laying period and then dechorionated 1-2 minutes in 50% bleach. Dechorionated embryos were transferred to a homemade plastic basket with a nylon mesh bottom, similar to reported previously (Rand et al., 2010). Embryos were then permeabilized in 1:10 Citrasolv (CitraSolv LLC, Danbury CT, USA) in water in a glass container with constant gentle shaking. Embryos were washed extensively with 1xPBS and then transferred to either a 10 mL hydroxyurea (HU) treatment solution (1xPBS, 100 uM Rhodamine B, 100 mM HU), or a 10 mL control solution (1xPBS, 100 uM Rhodamine B). Rhodamine B was included in these incubation solutions to mark embryos that had been successfully permeabilized. Embryos were incubated in the HU treatment or control solution for 40 minutes to maximize the number of interphase NC12 and NC13 embryos following fixation. In the case of the HU-treatment, the 40 min-duration also maximizes the number of DNA replication-inhibited NC12/NC13 embryos. The basket was removed from the incubation solution and blotted on a paper towel before the nylon mesh was removed. The embryos were immediately transferred to a glass scintillation vial filled with a formaldehyde fixation solution (2 mL 1xPBS/0.5% Triton X-100, 6 mL heptane, 180 ul 20% formaldehyde) by dunking the nylon mesh carrying the embryos in the vial, with the transfer taking ∼30 seconds. The embryo-free mesh was then removed from the vial and a standard embryo fixation protocol for ChIP-seq performed.

#### Embryo collection

For the collection of DNA replication-inhibited and control-treated embryos, following treatment and fixation embryos were sorted for ChIP-seq on 1%PBS-agarose plates in 1xPBS/0.5% TritonX-100. Sorting was performed using a Leica M165 FC Fluorescent Stereo Microscope to illuminate embryos with 488 nm light. NC12 and NC13 embryos were identified based on nuclear density indicated by Rpa70-GFP nuclear signal. During the collection of HU-treated embryos, embryos with most likely inhibited DNA replication were identified based on the distribution of Rpa70-GFP signal within nuclei. Embryos displaying punctate Rpa70-GFP within nuclei were preferentially selected, given the prior observation that replication-inhibition alters the intra-nucleus distribution of Rpa70-GFP (Blythe & Wieschaus, 2015b). 100 NC12/NC13 embryos were collected for each ChIP-seq replicate and stored at -80°C in 1xPBS/0.5% TritonX-100.

For the collection of *bcd osk tsl* and control *w* embryos, no treatment was performed. Embryos were collected after a 4-hr laying period, dechorionated, and fixed for ChIP-seq according to a standard protocol (Blythe & Wieschaus, 2015b). Fixed embryos were hand-sorted for NC14-stage embryos on 1%PBS-agarose plates in 1xPBS/0.5% TritonX-100. Sorting was performed under white light illumination by a Leica M165 FC Fluorescent Stereo Microscope. 50 NC14 embryos were selected for each ChIP-seq replicate based on morphological indicators including signs of cellularization and the clearing of lipid vesicles from the embryo’s periphery. Collected embryos were stored at -80°C in 1xPBS/0.5% TritonX-100.

#### Chromatin immunoprecipitation

Chromatin immunoprecipitations were performed as described previously (Blythe & Wieschaus, 2015b), with the following alterations. For each experiment, the embryos used for immunoprecipitation (IP) were 100 hand-sorted NC12/NC13 HU-treated Rpa70-GFP embryos and 100 hand-sorted NC12/NC13 control-treated Rpa70-GFP embryos, or 50 hand-sorted NC14 *bcd osk tsl* embryos and 50 hand-sorted NC14 *w* embryos. Each genotype was sonicated 4x 15 seconds at 20% Output and full duty cycle (Branson Sonifier 450) and then evenly split up into samples for the immunoprecipitations. ChIP was performed using a rabbit α-Bcd antibody (gift from J. Zeitlinger), a mouse α-Rpb1 CTD (4H8) monoclonal antibody (Cell Signaling Technology #2629), and a rabbit α-c-myc polyclonal antibody (Sigma-Aldrich #C3956) on HU-treated and control-treated embryos, with 25 embryo’s-worth of DNA in each IP. ChIP was performed using a mouse α-RNA Polymerase II CTD repeat YSPTSPS antibody (abcam #ab5408) and a rabbit α-c-myc polyclonal antibody (Sigma-Aldrich #C3956) on *bcd osk tsl* and *w* embryos, with 25 embryo’s-worth of DNA in each IP. Following incubation of the samples with blocked Protein G Dynabeads (Thermo Fisher), samples were washed twice in 10 mM Tris pH7.5 5 mM MgCl_2_ and then tagmented by exposure to 1 μl Nextera Tn5 Transposase (Illumina) while shaking at 1,000 rpm for 40 minutes at 37°C. All washes were performed after tagmentation as described in Blythe and Wieschaus, 2015 (Blythe & Wieschaus, 2015b). Following crosslink reversal and DNA cleanup, libraries were amplified as described in Blythe and Wieschaus, 2016 (Blythe & Wieschaus, 2016).

### QUANTIFICATION AND STATISTICAL ANALYSIS

### Modeling

Model structures, simulation method, and parameter sweeps are described in detail in the Supplemental Information.

### Enhancer sequence analysis

#### Predicting binding sites

PWM scores were calculated for each bp in the *hb* P2 sequence using the R Biostrings package (https://github.com/Bioconductor/Biostrings) and normalized by their maximum possible score. The Bcd and Zld PWMs were obtained from the UMass Chan Medical School Database of *Drosophila* TF binding specificities.

#### Predicting nucleosome positioning

The positions of likely nucleosomes were determined for the *hb* P2 sequence using NuPoP software (Xi et al., 2010) and identified at nucleotide positions 2-148 and 157-303 within a genomic sequence containing *hb* P2 flanked by 30 bp on its 5’ and 3’ ends. Similar positions were obtained by varying the flanking length of DNA for the sequence given to the NuPoP algorithm.

### Live-imaging analysis: MS2

All analyses of live-imaging data were performed in MATLAB (https://www.mathworks.com/products/matlab.html, R2022a). Live-imaging data was quantified during NC13, defined as the time period between the start of anaphase 12 and the start of anaphase 13.

#### Nuclei segmentation

Nuclei were segmented based on the His2Av-RFP channel. On a per-frame basis, the His2Av-RFP stack was filtered with a 3-D Guassian smoothing kernel with a standard deviation of 2 pixels. The smoothed stack was then binarized using Otsu’s method. Following binarization, morphological opening was performed using a spherical structuring element with a 4-pixel radius. Connected components with a radius of fewer than 5 pixels were then removed from the binary stack. Following an extended minima transformation, the stack was segmented with the watershed algorithm.

#### MS2 foci detection

To identify transcriptional foci, the MCP-GFP channel was processed on a per-frame basis. GFP intensity was first contrast-enhanced by employing top-hat and bottom-hat filtration (using a spherical structuring element with a 3-pixel radius). Difference of Gaussian (DoG) filtration was then performed on the contrast-enhanced stack, with 3-D Guassian smoothing kernels of 1-pixel and 5-pixel standard deviations. Pixels outside of the nuclear mask or with values below the estimated noise floor were then eliminated before the stack was binarized. Foci were identified from the binarized stack and eliminated if they had a radius of greater than 100 pixels or if they occupied fewer than 3 z-planes. Following segmentation of the DoG image with the binary mask, foci were eliminated if their total intensity was less than 100 AU.

#### AP position mapping

Per embryo, to determine the AP position of the imaged ROI, the normalized cross-correlation between a max-projection of the His2Av-RFP ROI during NC14 and a max-projection of the His2Av-RFP NC14 embryo overview image was calculated. The AP positioning of the ROI was identified by the location of highest correlation between the images.

#### Nuclei tracking

Nuclei were tracked frame-to-frame based on the segmentation of the His2Av-RFP channel. Centroids were calculated for all segmented nuclei in a frame and nuclei tracked by finding their centroid’s nearest neighbor in the following frame. If nuclei were lost track of during the time of active transcription, they were excluded from subsequent analyses.

#### Calculation of transcriptional features

The MS2 transcriptional dynamics of each nucleus over NC13 was calculated by segmenting the MCP-GFP channel with the 3-D MS2 foci mask and summing the fluorescence within each spot volume. To account for differences in nuclear cycle length between replicates, frames were sequentially divided into 100 bins—approximately the number of frames in the shortest NC13 of the MS2 imaging datasets—and the mean MS2 fluorescence for each nucleus calculated within each bin. Following binning, the nuclear fluorescence tracks were filtered to remove spots detected for durations less than the time required for transcription of the *y* intron of the reporter containing the MS2 stem loops (∼1.6 min given the prior measurement of a ∼2.5 kb/min Pol II elongation rate in NC13 (Fukaya et al., 2017)). Per-nucleus transcriptional onset times were calculated by finding the earliest bin in which each spot appeared following mitosis and converting the bin numbers into minutes given a 10-second frame time. Relative Pol II loading rates (AU/min) were calculated for each nucleus as the difference in a rolling average of spot fluorescence between the onset time of the spot and the time 1 minute after transcriptional onset. The fraction of active nuclei was calculated within 1% AP bins by dividing the number of nuclei that initiated transcription in a 1% AP span by the total number of nuclei measured within the bin.

### Live-imaging analysis: EGFP-Bcd

Nuclei were segmented and tracked over NC13 with the methods described in the MS2 analysis section. The GFP-Bcd channel was segmented using the nuclear mask, and the mean GFP fluorescence calculated within each nucleus over NC13.

### Fixed imaging analysis: HCR

For the Kr mutant experiment, *hbP2-MS2 Kr^1^/hbP2-MS2 Kr^1^* and control embryos were distinguished based on the Kr channel, as the *Kr^1^* mutant is RNA-null. For visualization, images were max-projected and the maximum pixel intensities set to allow for comparison of the posterior boundary of reporter expression between genotypes. As *hbP2-MS2 Kr^1^/SM6* (control) embryos were heterozygous for the reporter, the maximum intensity threshold of the 488 nm channel was lowered farther than that of the mutant embryos homozygous for *hbP2-MS2*. For the uniform Bcd experiment, images were max-projected and the maximum intensity threshold of the 488 nm channel set to be equivalent between control (*hbP2-MS2*/+) and mutant (*hbP2-MS2*/+ in uniform Bcd) embryos.

### ChIP-seq data analysis

Unique dual-indexed ChIP-seq libraries were subjected to paired-end 150 bp sequencing on an Illumina Novaseq X sequencer (Admera Health). Barcode-split fastq files were first trimmed of adapter sequences using TrimGalore (https://github.com/FelixKrueger/TrimGalore.git). Trimmed reads were mapped to the *Drosophila melanogaster* dm6 reference genome using Bowtie2 (v2.4.1) (Langmead & Salzberg, 2012) with option -X 2000. Sorted mapped reads were passed through Picard MarkDuplicates (v2.21.4, http://broadinstitute.github.io/picard/) to mark suspected duplicate reads. Non-duplicated mapped reads with map quality scores ≥ 10 were merged by replicate following import to R (v4.2.2) as a Genomic Ranges object using the GenomicAlignments package (Lawrence et al., 2013). To visualize coverage of ChIP-seq experiments, the average coverage of reads per 10 bp window was calculated and normalized to the total number of reads per merged sample/one million to yield counts-per-million reads per 10 bp window. Such coverage data were plotted using the GViz (v1.42.0) (Hahne & Ivanek, 2016).

## Author Contributions

**E.A.D.:** original draft preparation (lead); review and editing (equal); conceptualization (equal); methodology (equal); investigation (lead); software (lead); formal analysis (lead); visualization (equal); data curation (equal). **C.C.:** investigation (supporting). **N.M.M.:** methodology (equal), review and editing (supporting). **S.A.B.:** supervision and administration (lead); funding acquisition (lead); conceptualization (equal); methodology (equal); review and editing (equal); visualization (equal); data curation (equal).

## Acknowledgments

We are grateful to members of the Blythe Lab, as well as Chris Petersen, Hernan Garcia and Eric Wieschaus for comments on the manuscript. We are thankful to Julia Zeitlinger, who provided us with an antibody for Bicoid, and Mustafa Mir, for helpful conversations. We thank Bloomington Drosophila Stock Center, Hernan Garcia, Chris Rushlow, Steve Small, Pinar Onal, and Eric Wieschaus for *Drosophila* stocks. Finally, we thank Flybase for providing an essential resource to the *Drosophila* community. E.A.D. and C.C. were supported by the Cellular and Molecular Basis of Disease training program (T32GM008061). E.A.D. is supported by a NSF GRFP fellowship (DGE-2234667) and is a Data Science Fellow at the Northwestern Institute on Complex Systems. N.M.M. and S.A.B. were supported by the National Institute for Mathematics and Theory in Biology (Simons Foundations award MP-TMPS-00005320 and National Science Foundation award DMS-2235451). Experiments were supported by the National Institutes of Health grant R01HD101563 to S.A.B.. S.A.B. is a Pew Scholar in the Biomedical Sciences.

## Notes

### Competing Interest Statement

The authors have declared no competing interest.

