## Supplemental Information for "Bicoid-nucleosome competition sets a concentration threshold for transcription constrained by genome replication"

In the main text, we describe the development of a model structure that can accurately predict *hbP2-MS2* onset times in both wildtype and conditions of altered nucleosome stability. The success of this model hinges on its incorporation of the following aspects of the *in vivo* environment: nuclear Bicoid (Bcd) import and export dynamics, the interplay between transcription factors (TFs) and chromatin at an enhancer, and the prevalence of DNA replication early in nuclear cycle 13 (NC13). Here, we elaborate on the development and implementation of this model structure.

### Table of Contents


**1. Stochastic simulations** **1**
- 1.1 Replication-independent 2
- 1.2 Nucleosome, replication-dependent 3

**2. Nuclear Bcd concentration** **4**

**3. Parameter sweeps** **6**

**4. Definitions of $k_{on}$** **6**
- 4.1 $V_{max}$ parameter 7
- 4.2 Open chromatin non-cooperative models 7
- 4.3 Open chromatin cooperative model 8
- 4.4 Nucleosome-dependent model 12

**5. DNA replication-dependent model** **15**
- 5.1 Model structure 15
- 5.2 $V_{max}$ evaluation 17
- 5.3 Evaluation of modeled inter-origin distances 20

**7. Parameter summary** **20**


### 1. Stochastic simulations

As $k_{on}$ depends on the nuclear Bcd concentration, to simulate promoter activity an algorithm must allow $k_{on}$ to vary over time. Therefore, we employed a recently developed discretized implementation of the Gillespie algorithm that updates transition rates at each timestep given instantaneous nuclear TF concentration (Zhao et al., 2024). In our implementation, $k_{on}$ is re-estimated after each timestep given the nuclear Bicoid concentration at that time. This algorithm uses a fixed timestep, which we define as $\Delta t$ = 10 seconds to reflect the frame time used in our live-imaging measurements. Simulations were run over the first 15 minutes of a 20-minute nuclear cycle 13, the time prior to nuclear membrane breakdown. To simulate when nuclei will initiate transcription of the reporter, we adapted the algorithm to account for the fact each nucleus contains two copies of the reporter following DNA replication. We defined a cumulative transition rate ($\theta$) that reflects the probability a nucleus becomes transcriptionally active for the reporter, i.e., that any promoter of the two *hbP2-MS2* copies within the nucleus transitions into the ON state.

#### 1.1 Replication-independent

For the replication-independent models, a distribution of onset times was simulated as follows:

initialize AP position as $x = 20\%$

**while** $20\% \geq x \leq 80\%$ **do**

select a nucleus at the AP position $x$

initialize time as $t = 0$ and the promoter of each reporter copy as OFF

**while** $t \leq 15$ minutes **do**

with the promoters OFF at time $t$, calculate the nuclear Bcd concentration with

$$[Bcd](x, t) = Bcd_0 * e^{\frac{-x}{\lambda}} * Bcd_{norm}(t)$$

calculate $k_{on}$ given the current nuclear Bcd concentration and choice of the below model structures (elaborated upon later in the Supplement)

Open Chromatin Linear Model:

$$k_{on} = V_{max} * \frac{[Bcd]}{[Bcd]_{max}}$$

Open Chromatin Michaelis-Menten Model:

$$k_{on} = V_{max} * \frac{[Bcd]}{K_m + [Bcd]}$$

Open Chromatin Cooperative Model:

$$k_{on} = V_{max} * \frac{[Bcd]^{n_H}}{K_d + [Bcd]^{n_H}}$$

Nucleosome-dependent Model:

$$k_{on} = V_{max}(Y_{dist}([Bcd]) * Y_{prox}([Bcd]))$$

calculate the cumulative transition rate $\theta = 2k_{on}$

draw a random waiting time $\tau$ from $\tau \sim \text{Exp}(\theta)$

increment time such that $t = t + \Delta t$

decide whether a promoter transitioned in the timestep

**if** $\tau \geq \Delta t$ **then** both promoters remained in the OFF state

**else if** $\tau < \Delta t$ **then** a promoter transitioned to the ON state

record the onset time of the nucleus as $t$

increment AP position such that $x = x + 0.05\%$ and select a new nucleus

**end if**

**end while**

**end while**

#### 1.2 Nucleosome, replication-dependent

For the replication-dependent model, a distribution of onset times was simulated as follows:

initialize AP position as $x = 20\%$

**while** $20\% \geq x \leq 80\%$ **do**

determine when the reporter's promoter will complete DNA replication by drawing an inter-origin distance $X$ and converting it to a replication time $t_{rep}$ with

$X \sim \Gamma(\alpha = 2\, origins,\ \beta = \frac{9.7\, kb}{2\, origins})$ and $t_{rep} = \frac{X\, kb}{5.3\, kb/min} + 3.75\, min$

initialize time as $t = 0$ and the promoter state as OFF

**while** $t \leq 15$ minutes **do**

with all promoters OFF at time $t$, calculate the nuclear Bcd concentration with

$$[Bcd](x,t) = Bcd_0 * e^{\frac{-x}{\lambda}} * Bcd_{norm}(t)$$

calculate $k_{on}$ given the current nuclear Bcd concentration and the nucleosome-dependent model structure

$$k_{on} = V_{max}(Y_{dist}([Bcd]) * Y_{prox}([Bcd]))$$

calculate the cumulative transition rate $\theta$ given the promoter's replication state

**if** $t \leq t_{rep}$ **then** $\theta = k_{on}$

**else if** $t > t_{rep}$ **then** $\theta = 2k_{on}$

**end if**

draw a random waiting time $\tau$ from $\tau \sim \mathrm{Exp}(\theta)$

increment time such that $t = t + \Delta t$

decide whether a promoter transitioned in the timestep

**if** $\tau \geq \Delta t$ **then** the promoter/s remained in the OFF state

**else if** $\tau < \Delta t$ **then** determine the onset time of the nucleus

**if** $t \leq t_{rep}$ **then**

record the onset time of the nucleus as $t_{rep}$

increment AP position such that $x = x + 0.05\%$ and select a new nucleus

**else if** $t > t_{rep}$ **then**

record the onset time of the nucleus as $t$

increment AP position such that $x = x + 0.05\%$ and select a new nucleus

**end if**

**end if**

**end while**

**end while**

### 2. Nuclear Bcd concentration

The two-state model of promoter activity relies on Bcd concentration as an input variable. Therefore, to implement the model, we estimated nuclear Bcd concentration dynamics over NC13 at different positions along the Bcd gradient. To determine average nuclear Bcd dynamics, we live-imaged embryos expressing EGFP-Bcd and calculated the normalized mean nuclear GFP-Bcd fluorescence across the embryo over NC13, $Bcd_{norm}$(t) (Fig. S5A). $Bcd_{norm}$(t) represents the normalized Bcd dynamics within any nucleus along the AP axis, as while the maximum Bcd concentration a nucleus experience varies exponentially across space, normalizing the concentration measurements by their maximums produces roughly equivalent temporal dynamics between nuclei (Fig. S5B & C). To convert normalized fluorescence dynamics to Bcd concentrations, we then scaled the normalized curve by the maximum Bcd concentration at each AP position calculated according to the Synthesis-Diffusion-Degradation (SDD) model for formation of the Bcd gradient (Drocco et al., 2011; Gregor, Wieschaus, et al., 2007; Grimm et al., 2010; Little et al., 2011). The SDD model allows the Bcd concentration at any fractional AP position $x$ to be calculated with the exponentially decaying function

$$[Bcd](x) = Bcd_0 * e^{\frac{-x}{\lambda}}, \quad \textbf{S1}$$

where $Bcd_0$ is the maximum amount of Bcd protein at the anterior tip of the embryo and $\lambda$ is the fractional length scale of the gradient calculated according to

$$\lambda = \frac{\sqrt{D\tau}}{L_e}. \quad \textbf{S2}$$

Here, $D$ is the diffusion constant of Bcd, $\tau$ is the lifetime of the protein, and $L_e$ is the average length of the embryo. To determine the Bcd concentration profile, we evaluated Equation S1 across the AP axis with $Bcd_0$ = 140 nM (Abu-Arish et al., 2010), $D$ = 3 μm$^2$/second (Durrieu et al., 2018), $\tau$ = 50 minutes (Drocco et al., 2011), and $L_e$ = 500 μm (Fig. S5D). $Bcd_0$ = 140 nM represents a lower bound for the anterior nuclear Bcd concentration based on published FCS measurements (Abu-Arish et al., 2010) and $D$ = 3 μm$^2$/second falls within a published range of best-fit parameters to the Bcd gradient (Durrieu et al., 2018). $L_e$ = 500 μm is widely accepted. Application of the SDD model allowed us to convert our normalized EGFP-Bcd fluorescence measurements into Bcd concentrations by evaluating

$$[Bcd](x,t) = Bcd_0 * e^{\frac{-x}{\lambda}} * Bcd_{norm}(t). \qquad \textbf{S3}$$

*[Bcd]*(x,t) provides the Bcd concentration dynamics at any AP position over NC13. The two-state model of transcriptional activation defined in this work draws upon these dynamics to estimate how the probability of Bcd binding to its motifs in *hb* P2 changes across space and time, and how these binding dynamics relate to transcriptional timing.

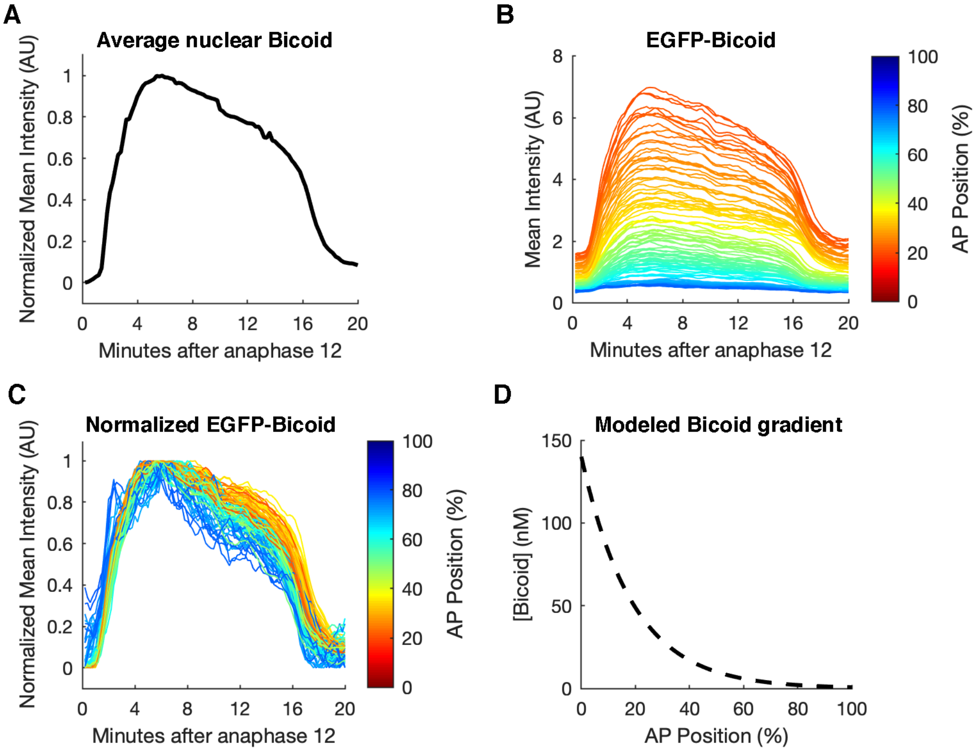


**Figure S5: Calculating nuclear Bicoid concentration dynamics.**

A) Normalized average nuclear Bicoid concentration dynamics. EGFP-Bicoid levels were averaged over NC13 for 218 nuclei across the AP axis. The background fluorescence was subtracted from the mean dynamics, and the dynamics then normalized by the maximum mean fluorescence.
B) Per-nucleus nuclear EGFP-Bicoid dynamics. Plotted are the nuclear concentration dynamics of 100 nuclei randomly chosen between 20-80% egg length. Intensity traces are colored according to the AP positions of the measured nuclei.
C) Normalized per-nucleus nuclear Bicoid dynamics. Plotted are the measurements in Panel B normalized by their maximums. Traces are colored according to the AP positions of the measured nuclei.

D) The Bicoid gradient estimated by the Sythesis-Diffusion-Degradation (SDD) model. The SDD model was applied using a maximum Bicoid concentration ($Bcd_O$) of 140 nM, a protein timetime ($\tau$) of 50 minutes, a diffusion coefficient ($D$) of 3 μm$^2$/second, and an embryo length ($L_e$) of 500 μm.

### 3. Parameter sweeps

To find best-fit parameters for different model structures, we evaluated the correspondence between model output and our *hbP2-MS2* fraction of active nuclei measurements (Fig. 2C). To summarize the output of a given parameter set, we fit a Hill equation, *f*, to the simulated fraction of active nuclei across Bcd concentrations, *[Bcd]*, specifying

$$f([Bcd]) = \frac{[Bcd]^{steepness}}{position^{steepness} + [Bcd]^{steepness}}. \quad \textbf{S4}$$

Here, *steepness* modulates the steepness of the posterior boundary of the fraction of active nuclei approximated by the Hill equation. *position* represents the Bcd concentration at which the fraction of active nuclei is half-maximal, therefore reflecting the position of the expression domain's posterior boundary. For a given combination of parameter values, we evaluated how well the *steepness* and *position* values of the Hill fit to the simulated fraction of active nuclei corresponded to the *steepness* and *position* of a Hill fit to the *hbP2-MS2* fraction of active nuclei (*steepness* = 8, *position* = 16.7 nM) (Fig. S6). In Figures S7, S8, S10, S11, and S12, we refer to the data's *steepness* value as 0.4 x/L, the AP position in fractional egg length where Bcd is expressed at 16.7 nM.

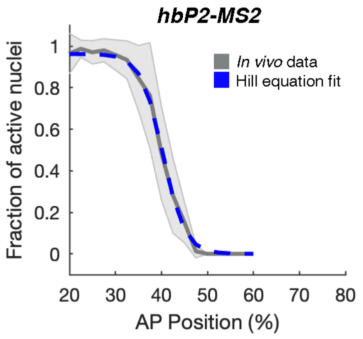


**Figure S6. Hill equation fit to the fraction nuclei active for *hbP2-MS2*.** Shown is the fraction of active nuclei plotted in Fig. 2C (gray) and the results of a Hill equation fit to the mean fraction of active nuclei (blue) with *steepness* = 8 and *position* = 16.7 nM (corresponding to an AP position of 40% egg length).

### 4. Definitions of $k_{on}$

$k_{on}$, the OFF-ON transition rate of the DNA replication-independent promoter model, depends on Bicoid concentration in space and time $k_{on} = V_{max}\, f([Bcd](x,t))$. We tested four different definitions of $k_{on}$ through varying the form of $f([Bcd](x,t))$ to determine the most likely mechanism by which Bcd activates transcription. As our ATAC-seq and ChIP-seq analyses indicate that Bcd competes with nucleosomes for occupancy at *hb* P2, we hypothesized that defining $k_{on}$ according to a TF-nucleosome competition mechanism would allow the model to most accurately simulate *hbP2-MS2* onset times (Fig. 1A & B). That said, we also ruled-out the possibility that Bcd interacts with its sites through alternative mechanisms, including noncooperative or pairwise cooperative binding to naked DNA. Here, we detail the application and results of changing the definition of $k_{on}$ in the two-state model of promoter activity. We analyze the following model structures: 1) two open chromatin, non-cooperative models (a Linear Model and a Michaelis Menten Model), 2) an open chromatin, cooperative model ("Hill Model"), and 3) a nucleosome-dependent model.

#### 4.1 $V_{max}$ parameter

In all model structures presented in this work $k_{on} = V_{max} f([Bcd])$, where the parameter $V_{max}$ represents the maximum possible rate at which a promoter can transition into the ON state and has units of transitions/10 seconds. We report the maximum number of transitions in 10 second intervals, as our live-imaging MS2 data was collected with a 10-second frame time. In order to estimate a value for $V_{max}$, we drew upon measurements of Bcd clustering dynamics within the nucleus. Recent measurements calculated the Bcd cluster lifetime as 2.4 ± 0.3 seconds and the inverse of cluster frequency as 1.6 ± 0.3 seconds (Munshi et al., 2024). In total, a Bcd cluster takes approximately 4 seconds to form and then disperse. Thus, in a 10 second timstep, 2.5 clusters can act at the *hb* P2 enhancer. We assume that the speed of cluster formation and dispersal sets the fastest rate at which a promoter can transition to the ON state. Therefore, given a 10 second time step, $V_{max}$ = 2.5 transitions/10 seconds. We view $V_{max}$ = 2.5 as an upper conceptual limit, given we assume an instantaneous promoter transition following Bcd binding to *hb* P2. In the main text, we report modeling results when $V_{max}$ = 1, a conservative estimate given the clustering measurements. Below we evaluate the impact of changing $V_{max}$ on the performance of the non-cooperative model structures, as well as on the best-fit parameters of the nucleosome, DNA replication-dependent model.

#### 4.2 Open chromatin non-cooperative models

We first tested a simple mechanism for Bcd's activity that could reflect how Bcd binding non-cooperatively to open chromatin impacts transcription. We defined $k_{on}$ simply as

$$k_{on} = V_{max} * \frac{[Bcd]}{[Bcd]_{max}}, \quad \textbf{S5}$$

where $k_{on}$ is linearly related to the normalized Bcd concentration. The parameter $V_{max}$ represents a maximum transition rate and is discussed above. Simulating a distribution of transcriptional onset times with this definition of $k_{on}$ and $V_{max}$ = 1 failed to recapitulate the measured *hbP2-MS2* onset time distribution (Fig. S7A). While the model does predict a correlation between mean onset time and AP position, it fails to predict a posterior boundary of MS2 expression, instead simulating expression across the entire AP axis (Fig. S7A). In an effort to find conditions that restrict the simulated expression to the high Bcd concentrations in the embryo's anterior, we performed a parameter sweep across $V_{max}$ = 0.01-1 in steps of 0.01. We find that, with a very small value of $V_{max}$ ($V_{max}$ = 0.04), the linear model can predict the *in vivo* data's *position* value of 0.4 x/L (Fig. S7B). However, the model cannot predict the *steepness* value of the *in vivo* data (*steepness* = 8) using any of the tested $V_{max}$ values (Fig. S7C). The results of this parameter sweep indicate that no value of $V_{max}$ can simultaneously predict both boundary position and steepness; while increasing $V_{max}$ slightly improves the prediction of boundary steepness, it dramatically reduces the accuracy of the boundary position prediction (Fig. S7B & C).

As a linear relationship between $k_{on}$ and Bcd concentration could not reproduce the *in vivo* data, we also tested whether nonlinearly relating $k_{on}$ to Bcd concentration could improve the model's predictions. As described in the main text, we defined $k_{on}$ according to Michaelis-Menten reaction kinetics, with

$$k_{on} = V_{max} * \frac{[Bcd]}{K_m + [Bcd]}. \quad \textbf{S6}$$

Here, $K_m$ is the Bcd concentration at which $k_{on}$ is half-maximal. Setting $K_m$ to 16.7 nM, the Bcd concentration at the boundary of the *hbP2-MS2* expression domain, does not allow the model to accurately predict onset times with $V_{max}$ = 1 (Fig. 4B, Fig. S7D). Again, the model simulates onset times across the entire AP axis, failing to predict the observed posterior boundary of expression around 45% egg length (EL). With parameter sweeps, we tested whether increasing the value of $K_m$ and decreasing the value of $V_{max}$ would improve simulation accuracy, as these changes decrease the probability of the OFF-ON transition. We evaluated the Michaelis-Menten model's output across a grid of parameter values spanning $V_{max}$ = 0.01-1 (sampled in steps of 0.01) and $K_m$ = 0-140 nM (sampled in steps of 10 nM). With $k_{on}$ defined according to Michaelis-Menten reaction kinetics, the model can simulate an accurate posterior boundary of expression with very small values of $V_{max}$ ($V_{max}$ = 0.01-0.06) (Fig. S7E). However, no tested parameter combination allowed the Michaelis-Menten model to accurately predict boundary steepness (Fig. S7F). Therefore, we pursued an alternative definition for $k_{on}$ that could allow the model to simulate a sharp posterior boundary.

#### 4.3 Open chromatin cooperative model

Prior studies indicate that Bcd binds cooperatively to *hb* P2 to produce the sharp posterior boundary of the anterior *hb* expression domain (Gregor, Tank, et al., 2007; Park et al., 2019). The Hill equation is often employed to model how binding levels change across ligand concentrations given pairwise cooperativity (Hill, 1913). Despite the utility of the Hill equation for explaining thresholding, a prior study has reported that Hill-like binding fails to sufficiently describe static *hb* expression measurements (Park et al., 2019). Therefore, we expected that incorporating pairwise-cooperative Bcd binding into the model would not enable prediction of the posterior boundary of MS2 expression, nor the timing of transcriptional onset. We systematically interrogated whether linearly relating $k_{on}$ to the Hill equation could allow the model to simulate the *hbP2-MS2* onset time and fraction of active nuclei measurements. In the two-state model, we specified

$$k_{on} = V_{max} * \frac{[Bcd]^{n_H}}{K_d + [Bcd]^{n_H}}, \quad \textbf{S7}$$

where $K_d$ is the dissociation constant of the simultaneous assembly of Bcd molecules on the enhancer and $n_H$ is the Hill coefficient, a parameter indicating the degree of Bcd binding cooperativity. $V_{max}$ is the maximum possible OFF-ON transition rate. We initially set $n_H$ = 9 to reflect the number of Bcd motifs identified in *hb* P2 (Fig. 1C). As there are no measurements of $K_d$ for the simultaneous binding of nine Bcd molecules to *hb* P2, we set $K_d = 16.7^9$ nM. With this definition of $K_d$, 16.7 nM represents the *EC50* of the equation, the Bcd concentration at which the transition rate is half-maximal. We deemed 16.7 nM an appropriate value for the *EC50*, as the *hbP2-MS2* fraction of active nuclei is half-maximal at 16.7 nM (40% EL) (Fig. S6). While with this parameter combination the model does simulate a posterior boundary of expression, the simulated boundary falls around 10% EL more posterior than the *in vivo* measurement of around 45% EL (Fig. S7G). Furthermore, it dramatically underestimates the mean and variance of onset times. To further characterize model performance, we evaluated the Hill-based model across a grid of parameter values spanning *EC50* = 10-40 nM (steps of 1 nM) and $n_H$ = 1-10 (steps of 1), with $V_{max}$ constant at 1. We find that when *EC50* > ~22 nM and $n_H$ > ~4, the model can accurately predict the data's *position* value of 0.4 x/L (Fig. S7H). When $n_H$ = 5 and $n_H$ = 6, the model produces its best predictions of the data's *steepness* value of 8 (Fig. S7I). While some of the tested parameter sets can predict both boundary steepness and position within 10% deviation of the *in vivo* data, no tested parameter set can simulate both boundary features within 5% deviation of the data's *steepness* and *position* values (Fig. S7J & K). Given the limited accuracy of the model's boundary predictions, as well as the infeasibility of an all-or-none binding pattern of nine independent Bcd molecules, we rule-out the possibility that Bcd regulates *hbP2-MS2* through pairwise cooperative binding to open chromatin (Brennan et al., 2023; Burz et al., 1998; Weiss, 1997).

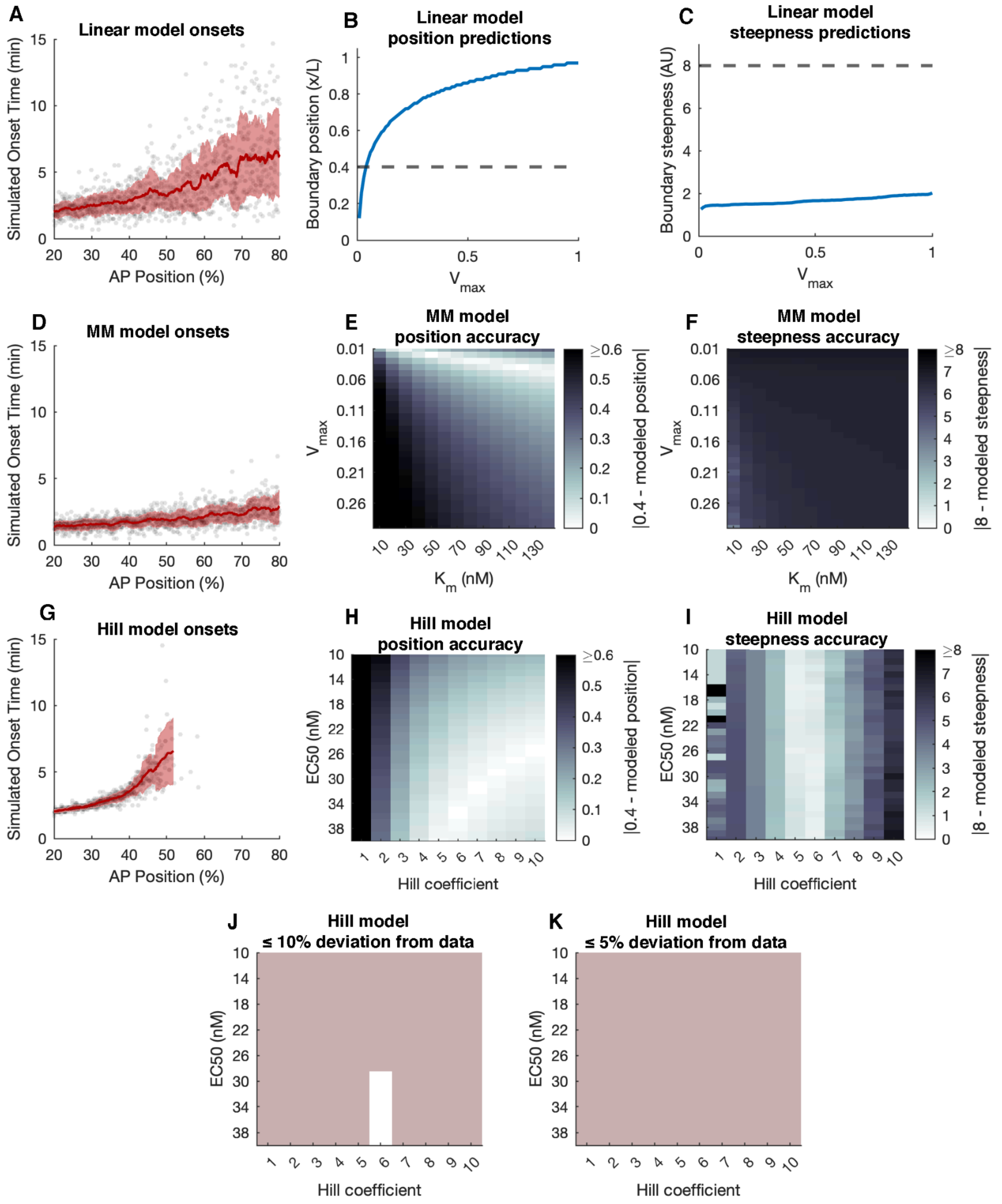


**Figure S7. Parameter sweep results for the open chromatin models.** Shown are onset time simulations and parameter sweep results for the Open Chromatin Linear Model, Open Chromatin Michaelis-Menten (MM) Model, and Open Chromatin Cooperative (Hill) Model. For all parameter sweeps, *position* and *steepness* values were determined by Hill fits to the simulated fraction of active nuclei. For each model structure, n = 4 parameter sweep replicates were performed, and the resulting *position* and *steepness* values averaged across replicates for each parameter combination. The average simulated *position* and *steepness* values are compared to the *position* and *steepness* values determined by a Hill fit to the *in vivo* fraction of active nuclei measurements (*position* = 0.4 x/L, *steepness* = 8).

A) Simulation output for the linear model. Shown is the result of a single simulation where $k_{on}$ is linearly related to the normalized nuclear Bicoid concentration and $V_{max}$ = 1. With this definition of $k_{on}$, the model fails to predict our *in vivo* measurements. The model predicts reporter transcription across the entire AP axis, as well as onset times substantially prior to the 4.5 minute onset time minimum observed *in vivo*.

B) Boundary position predictions from a parameter sweep for $V_{max}$ in the linear model. Shown are the *position* values (blue) determined from simulations performed with $V_{max}$ from 0.01 to 1 in steps of 0.01. The *in vivo position* value (*position* = 0.4 x/L) is indicated by the gray dashed line. A very small $V_{max}$ value ($V_{max}$ = 0.04) allows for prediction of the data's *position* value.

C) Boundary steepness predictions from a parameter sweep for $V_{max}$ in the linear model. Shown are the *steepness* values (blue) determined from simulations performed with $V_{max}$ from 0.01 to 1 in steps of 0.01. The *in vivo steepness* value (*steepness* = 8) is indicated by the gray dashed line. All tested values of $V_{max}$ substantially underestimate boundary steepness. The very slight upward trend of the steepness prediction suggests that a very high value of $V_{max}$ could predict boundary steepness, however, this improved steepness prediction would come at the cost of further reducing the accuracy of the simulated boundary *position* value (as indicated by Panel B).

D) Simulation output for the Michaelis-Menten (MM) model. Shown is the result of a single simulation where $k_{\mathrm{on}}$ is defined according to Michaelis-Menten reaction kinetics. The simulation was performed with parameter values of $V_{max}$ = 1 and $K_m$ = 16.7 nM. Like the linear model, the MM model fails to predict a posterior boundary of reporter expression, instead simulating transcriptional onset across the entire AP axis. Furthermore, this model systematically underestimates transcriptional onset times, frequently simulating onset times prior to the 4.5-minute minimum observed *in vivo*.

E) Boundary position predictions from parameter sweeps for $V_{max}$ and $K_m$ in the Michaelis-Menten model. Shown is a heatmap of the absolute value of the difference between the *position* values estimated from simulations and the *position* value of the *in vivo* data (*position* = 0.4 x/L). Simulations were performed across a grid of parameter values spanning $V_{max}$ = 0.01-1 (steps of 0.01) and $K_m$ = 0-140 nM (steps of 10 nM). Very small values of $V_{max}$ are required for accurate position estimates.

F) Boundary steepness predictions from parameter sweeps for $V_{max}$ and $K_m$ in the Michaelis-Menten model. Shown is a heatmap of the absolute value of the difference between the *steepness* values estimated from simulations and the *steepness* value of the *in vivo* data (*steepness* = 8). The same grid of parameter values is presented as in Panel E. No tested combination of $V_{max}$ and $K_m$ produces a *steepness* value that matches the *in vivo* data.

G) Simulation output for the Hill model. Shown is the result of a single simulation where $k_{on}$ is defined according to the Hill equation of cooperative ligand binding. The simulation was performed with parameter values of $V_{max}$ = 1, $n_H$ = 9, and *EC50* = 16.7 nM. The model simulates a posterior boundary of expression, yet substantially underestimates the mean and variance of onset times.

H) Boundary position predictions from parameter sweeps for *EC50* and $n_H$ in the Hill model. As in Panel E, shown is a heatmap of the absolute value of the difference between the *position* values estimated simulations and the *position* value of the *in vivo* data (*position* = 0.4 x/L). Simulations were performed across a grid of parameter values spanning *EC50* = 10-40 nM (steps of 1 nM) and $n_H$ = 1-10 (steps of 1). $V_{max}$ = 1 for all simulations. The best *position* predictions are made when $n > \sim 4$ and $EC50 > \sim 20$ nM.

I) Boundary steepness predictions from parameter sweeps for *EC50* and $n_H$ in the Hill model. As in Panel F, shown is a heatmap of the absolute value of the difference between the *steepness* values estimated from simulations and the *steepness* value estimated from the *in vivo* data (*steepness* = 8). The same grid of parameter values is presented as in Panel H. The best *steepness* predictions are made when $n_H$ = 5 or 6.

J) Parameter combinations that allow the Hill model to predict boundary *steepness* and *position* both within 10% of their *in vivo* values. Shown is a binary heatmap where white indicates a successful parameter combination, and gray indicates an unsuccessful parameter combination. With $n_H$ = 6 and $EC50 \geq 30$ nM, the Hill-based model can predict the data's *steepness* and *position* values within 10% deviation.

K) Parameter combinations that allow the Hill model to predict boundary *steepness* and *position* both within 5% of their *in vivo* values. Shown is a binary heatmap where white indicates a successful parameter combination, and gray indicates an unsuccessful parameter combination. No parameter combinations allow the Hill-based model to accurately predict the *in vivo* data.

#### 4.4 Nucleosome-dependent model

As detailed in the main text, Bcd competes with nucleosomes for occupancy at the hbP2 enhancer (Fig. 1A & B). In addition to the experimental measurements of Bcd binding and chromatin accessibility, the failure of the open-chromatin models points to the need for $k_{on}$ to take Bcd's interaction with nucleosomes into account. Also detailed in the main text, we drew upon a published mathematical description of TF-nucleosome competition (Mirny, 2010). This TF-nucleosome competition model emulates the Monod-Wyman-Changeux model of cooperative oxygen binding by hemoglobin (Monod et al., 1965). In the nucleosome competition case, a DNA locus can either be in an open state where Bcd can bind with a dissociation constant $K_O$, or in a nucleosomal state where Bcd can bind with a dissociation constant $K_N$. Given this allostery, the fractional occupancy of Bcd, $Y$, at $n$ Bcd motifs in a locus can be calculated with

$$Y = \frac{[Bcd]}{K_O} \frac{\left(1+\frac{[Bcd]}{K_O}\right)^{n-1} + L\frac{K_O}{K_N}\left(1+\frac{[Bcd]}{K_N}\right)^{n-1}}{\left(1+\frac{[Bcd]}{K_O}\right)^{n} + L\left(1+\frac{[Bcd]}{K_N}\right)^{n}}. \quad \textbf{S8}$$

In this equation, $L$ represents the equilibrium nucleosome occupancy in the absence of any bound TFs, and *[Bcd]* is the instantaneous Bcd concentration calculated as discussed above in the Supplemental Information. Both ATAC-seq data and NuPoP software (Xi et al., 2010) indicate two nucleosomes occupy *hb* P2. Therefore, we modeled Bcd's competition with each nucleosome separately as discussed in the main text. We defined $k_{on}$ as linearly related to the product of Bcd's fractional occupancy at the distal ($Y_{dist}$) and proximal nucleosomes ($Y_{prox}$), applying

$$k_{on} = V_{max}(Y_{dist} * Y_{prox}). \quad \textbf{S9}$$

With this definition of $k_{on}$, Bcd must occupy at least one of its sites at both the distal and proximal nucleosomes for the promoter to have the possibility to transition into the ON state.

Incorporating Bcd-nucleosome competition into the promoter activity model allows it to accurately predict the posterior boundary of *hbP2-MS2* expression with reasonable values of biophysical parameters describing the hbP2 enhancer sequence, as discussed in the main text (Fig. 4D). We identified best-fit parameter sets for the nucleosome-dependent model by performing a three-way parameter sweep across $K_O$ = 1-30 (steps of 1 nM), $K_N$ = 10-2810 (steps of 200 nM), and $L$ = 100-1000 (steps of 100), with $V_{max}$ = 1. We considered a range of $K_O$ values on the order of reported dissociation constants for Bcd's interaction with its binding sites as measured *in vitro* (Burz et al., 1998; Hannon et al., 2017). We constrained $L$ to the range of possible values inferred from experiments by Mirny, 2010 (Mirny, 2010). We computed the $K_N$ range from the $K_O$ range by instating the requirement that frequently $K_O/K_N < 0.1$, a ratio approximated by Mirny to lead to cooperative TF binding at a locus (Mirny, 2010). We held $V_{max}$ at 1, a lower bound estimate given Bcd clustering dynamics (Munshi et al., 2024). We find that, across the tested parameter sets, $K_O$ limits the number of parameter sets that can predict the position and steepness of the data's posterior boundary within 5% deviation from the *in vivo* measurements (Fig. S8A & B). Conversely, a wide range of $K_N$ values allows for accurate predictions of both boundary position and steepness, and a wide range of $L$ values allows for accurate prediction of boundary positions (Fig. S8A & B). However, $L$ values must be relatively high for the model to accurately predict boundary steepness (Fig. S8A & B). Considering the position and steepness predictions together, a total of 18 of the tested parameter sets allowed the model to predict both features of the boundary of the fraction of active nuclei within 5% deviation from the *position* and *steepness* values of the *in vivo* measurement (Fig. S8C). Out of these parameter sets, 9 predict the *in vivo* data's *position* value within 1% deviation from 0.4 x/L. We focus on $K_O$ = 9 nM, $K_N$ = 2610 nM, and $L$ = 700 in the main text and in the Supplemental Information, as it is included in these 9 parameter sets and closely matches the best-fit parameter set of the nucleosome, replication-dependent model discussed below ($K_O$ = 9 nM, $K_N$ = 2810 nM, and $L$ = 600).

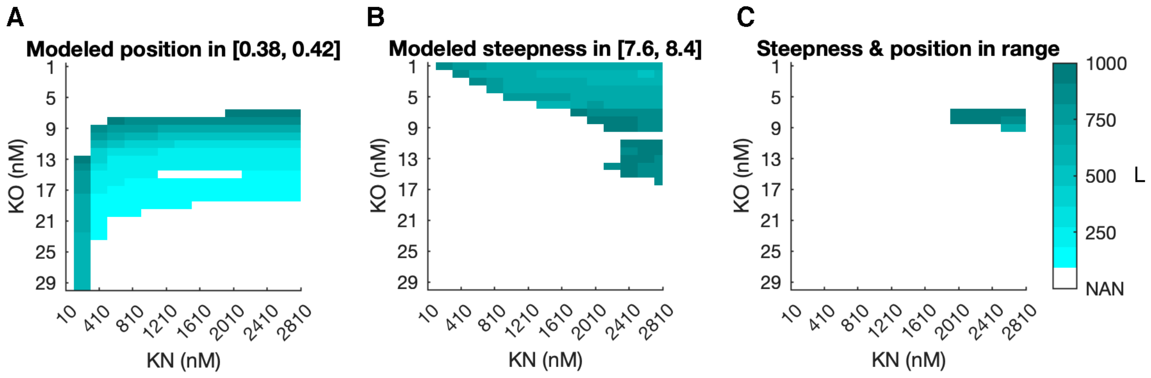


**Figure S8. Nucleosome-dependent model parameter sweep results.** Panels A-C show best-fit parameter sets for the nucleosome-dependent model determined by n = 4 replicates of a gridded parameter sweep spanning $K_O$ = 1-30 (steps of 1 nM), $K_N$ = 10-2810 (steps of 200 nM), and $L$ = 100-1000 (steps of 100), with $V_{max}$ = 1. Shown are heatmaps of $L$ values that, in combination with different $K_O$-$K_N$ pairs, allow for accurate prediction of boundary position (Panel A), boundary steepness (Panel B), or both boundary position and steepness (Panel C). White coloring at a $K_O$-$K_N$ combination indicates that, at that combination of $K_O$ and $K_N$ values, no $L$ value allows for accurate simulation of the specified boundary feature(s). The mean $L$ value across parameter sweep replicates is shown in cyan. Heatmaps indicate parameter combinations that predict the measured posterior boundary **(A)** position within 5% deviation of *position* = 0.4 x/L, **(B)** steepness within 5% deviation of *steepness* = 8, and **(C)** steepness and position within 5% deviation of their target values. In total, 18 of the tested parameter sets led to accurate boundary predictions.

Despite its ability to accurately predict the posterior boundary of the *hbP2-MS2* fraction of active nuclei, this nucleosome-dependent model fails to predict adequately delayed and variable transcriptional onset times in the embryo's anterior with $K_O$ = 9 nM, $K_N$ = 2610 nM, and $L$ = 700 (Fig. 4D). We sought to test whether increasing $K_O$, $K_N$, or $L$, could increase the mean and variance of the model's onset time predictions at high Bcd concentrations. Decreased Bcd-DNA binding affinities and increased nucleosome stability could lead to delays in transcriptional onset. We find that increasing $K_O$ from 9 nM to 14 nM in the baseline parameter set ($K_O$ = 9 nM, $K_N$ = 2610 nM, $L$ = 700) does not allow the model to fully reproduce the *in vivo* onset measurements (Fig. S9A & B). Furthermore, this increase in $K_O$ leads to an anterior shift in the simulated expression, reducing the accuracy of the modeled boundary prediction (Fig. S9A). Increasing $K_N$ from 2610 nM to 5000 nM in the baseline parameter set also does not increase the accuracy of the model's predictions; the model still simulates onset times in the anterior 20-30% EL prior to the 4.5-minute *in vivo* onset time minimum (Fig. S9C & D). Finally, increasing $L$ from 700 to its estimated maximum value of 1000 (Mirny, 2010) also does not improve the onset time predictions in 20-30% EL, with onset times predicted substantially prior to the *in vivo* measurements (Fig. S9E & F). $L$ = 1000 also shifts the posterior boundary prediction towards the anterior and away from the *in vivo* expression limit of 45% EL (Fig. S9E). We needed to add an additional feature to the model's structure—the timing of DNA replication—to increase the mean and variance of onset time predictions in the embryo's anterior without disrupting the posterior boundary prediction.

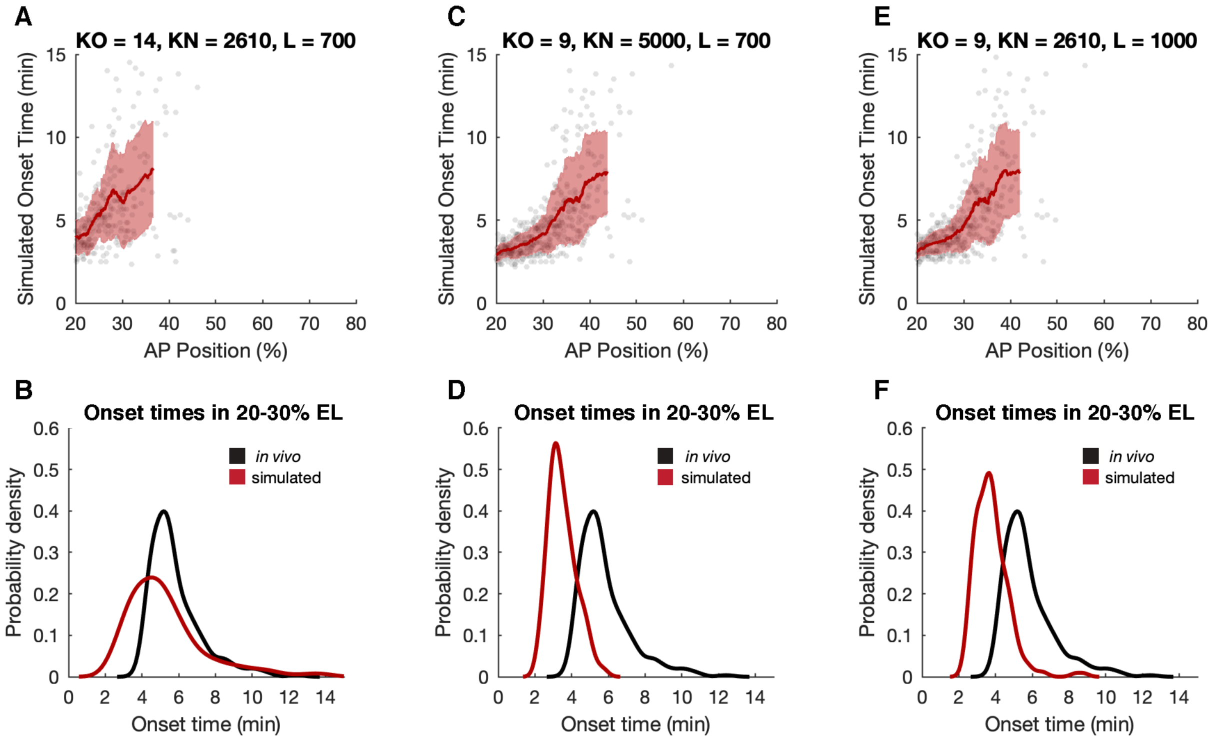


**Figure S9. Premature anterior onset time predictions by the nucleosome-dependent model.** Panels A, C, and E show the results of single onset time simulations by the nucleosome-dependent model using the specified parameter sets. Panels B, D, and F show kernel density estimates for the simulated onset times in 20-30% egg length in the simulations presented in the panels above (red), as well as the kernel density estimate for the measured *in vivo* onset times in 20-30% egg length (black). Simulations were run after changing single parameter values in the baseline parameter set of $K_O = 9$, $K_N = 2610$, and $L = 700$.

A) Increasing $K_O$ in the nucleosome-dependent model does not adequately delay onset times in the far anterior. Compared with the results for the baseline parameter set presented in Fig. 4D, while increasing $K_O$ from 9 to 14 nM shifts the posterior boundary towards the anterior, some onset times are still simulated prior to 4.5 minutes in the far anterior.

B) When $K_O$ = 14, $K_N$ = 2610, and $L$ = 700, simulated onset times in the anterior undershoot the *in vivo* measurements.

C) Increasing $K_N$ in the nucleosome-dependent model does not delay onset times in the anterior. Increasing $K_N$ from 2610 to 5000 nM does not noticeably change the simulated onset time distribution, as compared to the simulation results for the baseline parameter set presented in Fig. 4D.

D) When $K_O$ = 9, $K_N$ = 5000, and $L$ = 700, simulated onset times in the anterior undershoot the *in vivo* measurements.

E) Increasing $L$ in the nucleosome-dependent model does not delay onset times in the anterior. Increasing $L$ from 700 to 1000 only slightly changes the simulated onset time distribution, as compared to the simulation results for the baseline parameter set presented in Fig. 4D.

F) When $K_O$ = 9, $K_N$ = 2610, and $L$ = 1000, simulated onset times in the anterior undershoot the *in vivo* measurements.

### 5. DNA replication-dependent model

#### 5.1 Model structure

While the model can predict the posterior boundary of *hbP2-MS2* expression with $k_{on}$ defined according to Bcd's competition with the nucleosomes at *hb* P2, it fails to accurately predict the timing of transcriptional onset in the far anterior of the embryo (Fig. 4D, Fig. S9A-F). Therefore, we incorporated transcrption's dependence on the dynamics of DNA replication into the model. As discussed in the main text, we estimated the distances between origins of replication with the following gamma distribution,

$$X \sim \Gamma(\alpha = 2\, origins,\ \beta = \frac{9.7\ kb}{2\ origins}). \qquad \textbf{S10}$$

Here, the inter-origin distance $X$ has units of kilobases. To convert $X$ to a time following mitosis at which the reporter's promoter would complete replication, $t_{rep}$, we divided the distances by the speed of DNA polymerase, 5.3 kilobases/minute, and accounted for the estimated ~3.75 minute delay before replication initiation following mitosis 12:

$$t_{rep} = \frac{X\ kb}{5.3\ kb/min} + 3.75\ min. \qquad \textbf{S11}$$

To incorporate the timing of the promoter's replication into the model, we instated the requirement that, for the promoter to transition into the ON state, it must complete DNA replication and Bcd must bind to the hbP2 enhancer (Fig. 6A). The model defines $k_{on}$ with Eq. S9, as in the DNA replication-independent, nucleosome competition model. As presented in the main text, this nucleosome and replication-dependent model structure can simulate both the mean and variance of our *in vivo* onset time measurements (Fig. 6D-F).

We evaluated how well the nucleosome, replication-dependent model can predict the posterior boundary of the *hbP2-MS2* fraction of active nuclei across the grid of parameter combinations used for the nucleosome-dependent model parameter sweep presented previously. The nucleosome, replication-dependent model accurately predicts boundary position and steepness using similar parameter values to the best-fit parameters of the nucleosome-dependent model (Fig. S8A-C, Fig. S10A-C). We find the nucleosome, replication-dependent model best predicts the *in vivo* fraction of active nuclei measurements—estimating the *position* and *steepness* parameters within 5% deviation of their *in vivo* values—when it defines nucleosome stability at *hb* P2 as at least moderately high ($L \geq 600$), Bcd's affinity for its binding sites in the nucleosomal state as relatively low ($K_N \geq$ 1,610 nM), and Bcd's affinity for sites in the open state as relatively high ($K_O$ = 7-9 nM) (Fig. S10C). As a narrow range of $K_O$ values results in good predictions, we conclude a dissociation constant of ~8 nM describes Bcd's binding to its sites in *hb* P2 *in vivo*, narrowing down prior *in vitro* estimates (Fig. S10C, Fig. S11A) (Burz et al., 1998;

Burz & Hanes, 2001; Hannon et al., 2017; Ma et al., 1996). Therefore, we propose the eviction of nucleosomes at the boundary of the *hb* P2-mediated expression domain requires a high-affinity interaction between Bcd and its binding sites. In contrast to the model's low tolerance for variability in $K_O$, the model simply requires that $K_N$ and $L$ pass thresholds for it to accurately predict the *in vivo* data (Fig. S10C). The best-fit parameter sets contain high $K_N$ and $L$ values (Fig. S10C, Fig. S11B & C). Therefore, Bcd likely has a very low affinity for nucleosomal DNA and nucleosomes likely stably occupy *hb* P2. Furthermore, the low $K_O/K_N$ ratio achieved by all of the best-fit parameters sets falls within the range of values specified by Mirny as required for cooperative TF binding (Mirny, 2010) (Fig. S11D). The presence of nucleosomes at *hb* P2 likely increases the cooperativity of Bcd binding, leading to the formation of a sharp posterior boundary of *hbP2-MS2* expression.

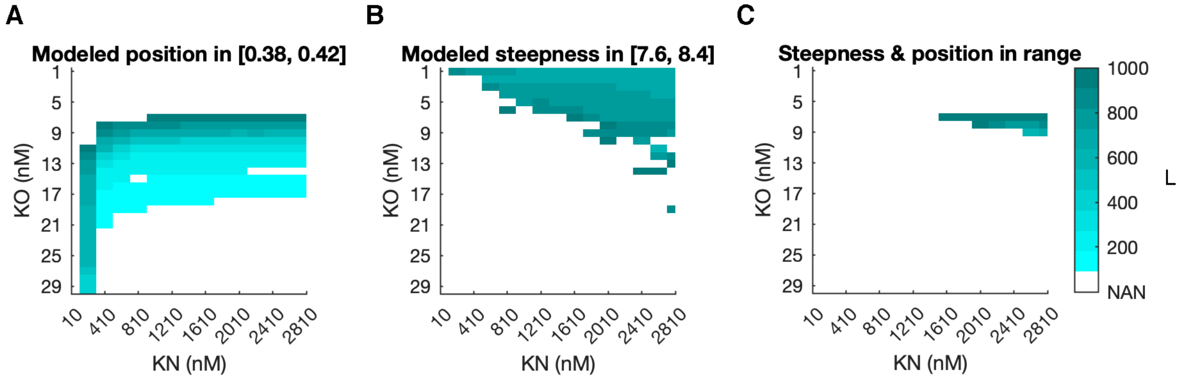


**Figure S10. Nucleosome and replication-dependent model parameter sweep results.** Panels A-C show best-fit parameter sets for the nucleosome, replication-dependent model determined by n = 4 replicates of a gridded parameter sweep spanning $K_O$ = 1-30 (steps of 1 nM), $K_N$ = 10-2810 (steps of 200 nM), and $L$ = 100-1000 (steps of 100), with $V_{max}$ = 1. Shown are heatmaps of $L$ values that, in combination with different $K_O$-$K_N$ pairs, allow for accurate prediction of boundary position (Panel A), boundary steepness (Panel B), or both boundary position and steepness (Panel C). White coloring at a $K_O$-$K_N$ combination indicates that, at that combination of $K_O$ and $K_N$ values, no $L$ value allows for accurate simulation of the specified boundary feature(s). The mean $L$ value across parameter sweep replicates is shown in cyan. Heatmaps indicate the parameter combinations in the nucleosome, replication-dependent model that predict the measured posterior boundary **(A)** position within 5% deviation of *position* = 0.4, **(B)** steepness within 5% deviation of *steepness* = 8, and **(C)** steepness and position within 5% deviation of their target values. In total, across parameter sweep replicates, 27 of the tested parameter sets led to accurate boundary positions. Results are similar to those reported for the replication-independent model in Fig. S8A-C.

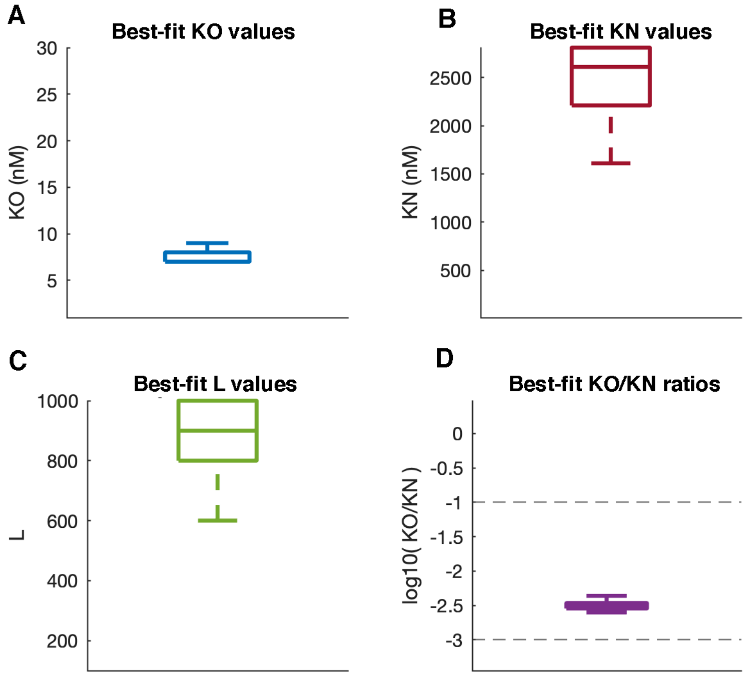


**Figure S11.** Distributions of best-fit parameter values for the nucleosome, replication-dependent model. Panels describe the values within the 27 parameter sets found in n = 4 parameter sweeps to lead to boundary *steepness* and *position* predictions within 5% of the data's boundary *steepness* = 8 and *position* = 0.4 x/L. In each parameter sweep replicate, parameters were varied across $K_O$ = 1-30 (steps of 1 nM), $K_N$ = 10-2810 (steps of 200 nM), and $L$ = 100-1000 (steps of 100), with $V_{max}$ = 1. Parameter sweep results for the nucleosome, replication-dependent model are also presented in Fig. S10.

A) A small range of $K_O$ values leads to accurate boundary predictions. Shown is the distribution of $K_O$ values within the best-fit parameter sets. The best-fit parameter sets contain a small range of $K_O$ values, 7-9 nM.
B) High $K_N$ values are required for accurate boundary predictions. Shown is the distribution of $K_N$ values within the best-fit parameter sets. The best-fit parameter sets contain relatively high $K_N$ values, 1610-2810 nM.
C) High $L$ values lead to accurate boundary predictions. Shown is the distribution of $L$ values within the best-fit parameter sets. $L$ values range from 600 to 1000, relatively high values in the $L$ range of 100-1000 specified in Mirny, 2010.
D) The best fit parameter sets contain the expected low $K_O/K_N$ ratio. Shown is a distribution of the $K_O/K_N$ ratios within each of the best-fit parameter sets. Gray dashed lines indicate the lower and upper bounds of $K_O/K_N$ designated by Mirny, 2010. All good parameter sets have a $K_O/K_N$ ratio that falls within the expected range of 0.1-0.001.

#### 5.2 $V_{max}$ evaluation

For our application of the open chromatin cooperative model and the nucleosome-dependent models, we set $V_{max}$ = 1, which we consider a conservative estimate for $V_{max}$ given prior measurements of Bcd clustering dynamics within the nucleus (Munshi et al., 2024). We wanted to explore how the value of $V_{max}$ impacts the best-fit values of $K_O$, $K_N$, and $L$. Therefore, we performed parameter sweeps at lower ($V_{max}$ = 0.01, 0.5) and higher ($V_{max}$ = 2.5) values of $V_{max}$. As discussed previously, given Bcd clusters take on average 4 seconds to form and then disperse, $V_{max}$ = 2.5 transitions/10 seconds represents an upper conceptual limit (Munshi et al., 2024). We find that, with $V_{max}$ = 0.01, few parameter sets allow the model to reproduce the observed boundary position or steepness (Fig. S12A & B). None of the 4500 tested parameter sets accurately predict both position and steepness when $V_{max}$ = 0.01 (Fig. S12C). With $V_{max}$ = 0.5, more parameter sets allow the model to reproduce the observed boundary position or steepness, and of the tested sets, 30 allow the model to predict both *position* = 0.4 and *steepness* = 8 within 5% deviation (Fig. S12D-F). Parameter sweeps when $V_{max}$ = 0.5 and $V_{max}$ = 1 yield very similar results, with 27 of the tested parameter sets accurately predicting both steepness and position when $V_{max}$ = 1 (Fig. S12D-I). When $V_{max}$ = 2.5, while parameter sets allow the model to make accurate position and steepness predictions independently, only 6 of the tested parameter sets lead to accurate prediction of both boundary position and steepness (Fig. S12J-L). $V_{max}$ = 2.5 approximates an upper bound beyond which no $K_O/K_N/L$ combination produces accurate boundary simulations.

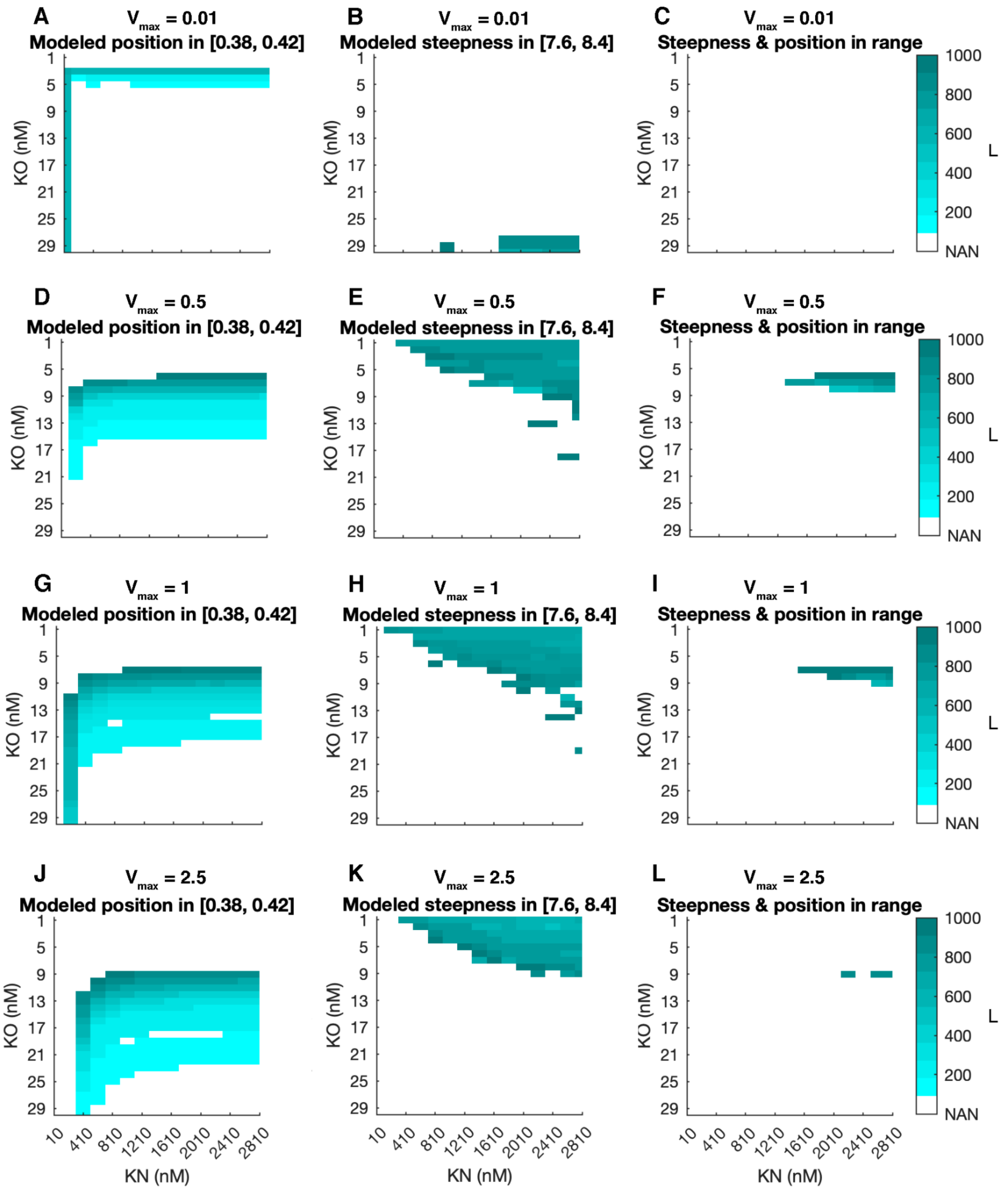


**Figure S12. The effects of changing the value of $V_{max}$ on the best-fit parameter sets of the nucleosome, replication-dependent model.** Panels A-L show best-fit parameter sets for the nucleosome, replication-dependent model with different values of $V_{max}$ ($V_{max}$ = 0.01, 0.5, 1, or 2.5) determined by gridded parameter sweeps spanning $K_O$ = 1-30 (steps of 1 nM), $K_N$ = 10-2810 (steps of 200 nM), and $L$ = 100-1000 (steps of 100). Shown are heatmaps of $L$ values that, in combination with different $K_O$-$K_N$ pairs, allow for accurate prediction of boundary position (Panels A, D, G, and J), boundary steepness (Panels B, E, H, and K), or both boundary position and steepness (Panels C, F, I, and L). White coloring at a $K_O$-$K_N$ combination indicates that, at that combination of $K_O$ and $K_N$ values, no $L$ value allows for simulation of the specified boundary feature(s) within 5% deviation of the data. Parameter sets are plotted in color when they can predict boundary position within 5% deviation of 0.4 x/L (in [0.38, 0.42] x/L), boundary steepness within 5% deviation of 8 (in [7.6, 8.4]), or both. The mean $L$ value across parameter sweep replicates (n = 4 per $V_{max}$ value) is shown in cyan. Heatmaps indicate parameter sets that allow for accurate prediction of boundary **(A)** position when $V_{max}$ = 0.01, **(B)** steepness was $V_{max}$ = 0.01, **(C)** position and steepness when $V_{max}$ = 0.01, **(D)** position when $V_{max}$ = 0.5, **(E)** steepness when $V_{max}$ = 0.5, **(F)** position and steepness when $V_{max}$ = 0.5, **(G)** position when $V_{max}$ = 1 (Fig. S10A), **(H)** steepness when $V_{max}$ = 1 (Fig. S10B), **(I)** position and steepness when $V_{max}$ = 1 (Fig. S10C), **(J)** position when $V_{max}$ = 2.5, **(K)** steepness when $V_{max}$ = 2.5, **(L)** position and steepness when $V_{max}$ = 2.5. In total, across parameter sweep replicates, 0, 30, 27, and 6 of the tested parameter sets allow the model to predict both position and steepness within 5% deviation of their target values when $V_{max}$ = 0.01, 0.5, 1, and 2.5 respectively.

#### 5.3 Evaluation of modeled inter-origin distances

To evaluate the goodness of fit of the gamma distribution (Eq. S10) to previously measured inter-origin distances reported (Blumenthal et al., 1974), we performed a KS test between the 1974 measurements and our gamma distributed modeled inter-origin distances. The previously measured inter-origin distances were reported in a histogram in Figure 4 of Blumenthal et al., with count data binned into 1 kb bins (1 to 30 kb), plus one bin for values greater than 30 kb. The bar heights reflected the fraction of total measurements (n = 316) with an observation within the binned kb value. We estimated the original 1974 count data by measuring histogram bar heights with a millimeter ruler and converting heights to fraction of measured distances by linear regression of the y-axis labels (0, 5, 10, and 15%) and their corresponding measured heights. We converted our reproduction of the 1974 data to counts per 1 kb bin by multiplying by the total number of measurements (316) and rounding to the nearest integer value. For these conversions, we dropped the > 30 kb bin since these data were not reported precisely. Our reproduction of the 1974 data is plotted in Figure 6B, purple bars. By eye, our estimate of the original data is accurate.

We performed $1 \times 10^3$ independent replicates of a KS test comparing our reproduction of the 1974 data with $1 \times 10^4$ randomly sampled values from the gamma distribution model. All $1 \times 10^3$ tests yielded p-values greater than 0.05, indicating that both samples are likely drawn from the same statistical distribution.

### 7. Parameter summary

| Parameter | Evaluated range | Reference |
|---|---|---|
| Maximum Bicoid concentration ($Bcd_0$) | 140 (nM) | Abu-Arish et al., 2010 |
| Bicoid diffusion constant ($D$) | 3 (μm$^2$/second) | Durrieu et al., 2018 |
| Bcd protein lifetime ($\tau$) | 50 (minutes) | Drocco et al., 2011 |
| Embryo length ($L_e$) | 500 (μm) | This paper |
| Maximum OFF-ON transition rate ($V_{max}$) | 0.01-2.5 (transitions/10 seconds) | This paper, based on Munshi et al., 2024 |

| | | |
|---|---|---|
| Dissociation constant of Bicoid binding to a single site in the Open Chromatin Michaelis-Menten Model ($K_m$) | 0-140 (nM) | This paper, based on Burz et al., 1998; Burz & Hanes, 2001; Hannon et al., 2017; Mat et al., 1996 |
| Bicoid concentration at which the OFF-ON transition rate is half-maximal in the Open Chromatin Cooperative Model (*EC50*) | 10-40 (nM) | This paper |
| Hill coefficient ($n_H$) | 1-10 (AU) | This paper |
| Bicoid's affinity for the open state in the nucleosome-dependent models ($K_O$) | 1-30 (nM) | This paper, based on Burz et al., 1998; Burz & Hanes, 2001; Hannon et al., 2017; Mat et al., 1996 |
| Bicoid's affinity for the nucleosomal state in the nucleosome-dependent models ($K_N$) | 10-2810 (nM) | This paper, based on Mirny, 2010 |
| Equilibrium between the open and nucleosomal states in the nucleosome-dependent models ($L$) | 10-1000 (AU) | Mirny, 2010 |
| Number of Bicoid binding sites in the nucleosome-dependent models ($n$) | proximal nucleosome: 4<br>distal nucleosome: 5 | This paper |
| Shape parameter in the gamma distribution of inter-origin distances ($\alpha$) | 2 (origins) | This paper, based on Blumenthal et al., 1974 |
| Scale parameter in the gamma distribution of inter-origin distances ($\beta$) | 9.7/2 (kb/origins) | This paper, based on Blumenthal et al., 1974 |

**Table S1: Parameter ranges used in the parameter sweeps.** Parameter ranges are reported for the parameter sweeps for the Open Chromatin Linear Model, Open Chromatin Michaelis-Menten Model, Open Chromatin Cooperative Model, Nucleosome-dependent Model, and Nucleosome/Replication-Dependent Model. Single values are reported for parameters held constant during the sweeps. References are supplied for parameter values directly specified in prior work, and for parameter values extrapolated from prior work.
